# Massively parallel interrogation of the fitness of natural variants in ancient signaling pathways reveals pervasive local adaptation

**DOI:** 10.1101/2024.10.30.621178

**Authors:** José Aguilar-Rodríguez, Jean C. C. Vila, Shi-An A. Chen, Manuel Razo-Mejia, Olivia M. Ghosh, Daniel F. Jarosz, Hunter B. Fraser, Dmitri A. Petrov

## Abstract

The nature of standing genetic variation remains a central debate in population genetics, with differing perspectives on whether common variants are almost always neutral as suggested by neutral and nearly neutral theories or whether they can commonly have large functional and fitness effects as proposed by the balance theory. We address this question by mapping the fitness effects of over 9,000 natural variants in the Ras/PKA and TOR/Sch9 pathways—key regulators of cell proliferation in eukaryotes—across four conditions in *Saccharomyces cerevisiae*. While most variants are neutral in our assay, ∼3,500 exhibited significant fitness effects. These non-neutral variants tend to be missense and to affect conserved, more densely packed, and less solvent-exposed protein regions. While some of these non-neutral variants are younger and rarer, and more often found in heterozygous states—consistent with purifying selection—a substantial fraction is present at high frequencies in the population, which is expected under balancing selection. Indeed, we find that variants with a positive fitness effect in our laboratory measurement show strong signs of local adaptation as they tend to be found specifically in domesticated strains isolated from human-made environments. Our findings support the view that while many common variants might be effectively neutral, a significant proportion have locally adaptive functional consequences and are driven into a subset of the population by local positive selection. This study highlights the potential to combine high-throughput precision genome editing with fitness measurements to explore natural genetic variation on a pathway-wide scale, thereby bridging the gap between population genetics and functional genomics to understand the nature of evolutionary forces in the wild.

## Introduction

One of the longstanding debates in population genetics centers on the nature of standing genetic variation (*1–3*). The classical theory (*4*), with nearly neutral theory being its latest incarnation (*5–9*), posits that standing genetic variation is generated through the balance between neutral and deleterious mutation and primarily shaped by purifying selection. This model predicts that while variants present at low population frequencies can be both neutral, nearly neutral, or even strongly deleterious, variants present at high frequencies are very unlikely to be deleterious and therefore must be either neutral or nearly neutral. This further implies that common variants are unlikely to have a substantial functional or fitness impacts. This perspective commonly serves as the null model for studies of genetic variation in the wild (*10*).

In contrast, balance theory argues that while purifying selection does keep deleterious alleles at lower frequencies, a meaningful proportion of common genetic variation is functionally important, beneficial in at least a subset of environments, and maintained at high frequencies through the action of balancing selection (*11*). Balancing selection can operate through many mechanisms, including overdominance (heterozygote advantage) (*12*, *13*), frequency-dependent selection (*14*), and selection that varies across space (local adaptation) (*15*) or time (fluctuating selection) (*16*).

Despite much effort and voluminous data, the current population genetic evidence remains inconclusive about which theory best describes population genetic variation (*2*, *3*, *17*). There is clear evidence for most genes that mutations that are more likely to be functional *a priori*, for instance, nonsynonymous variants—those that alter the amino acid sequence of proteins—tend to be present at lower allele frequencies than synonymous variants (*18–24*). This implies that they are likely to be disproportionately deleterious. While this pattern is consistent with the classical nearly neutral theory, in the absence of experimental functional data, it is hard to know whether all observed common nonsynonymous or other functional variants are neutral or whether they do contain a substantial fraction of balanced, functionally, and selectively consequential variants.

In principle it should be possible to resolve the debate about the functional consequences of the common variation by making direct measurements of the functional significance of individual common variants. However, quantitative genetics approaches, such as quantitative trait loci (QTL) mapping or genome-wide association studies (GWAS), almost universally lack the spatial resolution necessary to determine the functional or fitness consequences of single-nucleotide variants, which are precisely the type of genetic polymorphism most commonly found in natural populations (*20*).

Recent advances in precision genome editing allow us to address this question experimentally. While older CRISPR editing methods suffered from low efficiencies (5-20%), limiting the feasibility of massively parallel editing, new techniques in the budding yeast *Saccharomyces cerevisiae* have substantially improved precision genome editing efficiencies (80-100%) (*25–30*). This breakthrough now enables the simultaneous introduction of tens of thousands of variants in parallel and the accurate measurement of their fitness *en masse* using genomic barcodes.

In this study we have employed one such method, CRISPEY-BAR+, to map the fitness effects of more than 9,000 natural genetic variants with high precision. We chose variants segregating within the core Ras/PKA and TOR/Sch9 pathways, which control cell proliferation in response to environmental stimuli across all eukaryotes (*31–33*). These pathways are crucial for the virulence of several pathogenic yeasts (*34*, *35*), and the pathogenesis of diverse human cancers (*36*), and are often mutated to adaptive states in experimental evolution (*37–42*). Both standing genetic variation and *de novo* mutations within them contribute to the adaptation to novel nutrient sources in yeast (*37–47*), indicating that they may act as a “hot-spot” of evolutionary novelty and possibly functional genetic variation segregating in this species. The variants we investigate encompass the majority of the known coding and non-coding genetic variation in these pathways across this species (*20*). We therefore built a comprehensive “fitness landscape” (*48*)—a map linking genetic variation to fitness variation—for the standing genetic variants in these pathways. The environments we chose for the interrogation of fitness effects are known to yield large fitness variation for mutations in these pathways (*42*, *46*).

In this study, we find that large-effect functional variants do tend to be disproportionately deleterious, although this relationship is weak. What is surprising and inconsistent with nearly neutral theory is that many very large-effect polymorphisms are present at high frequencies in the population. Moreover, we find that variants that have positive fitness effects in our laboratory measurements are enriched in domesticated and industrial strains, suggesting that they have been driven to high frequency by local positive selection during adaptation to fermented food and other human-made environments (*49*). These results provide strong support in favor of the balance theory of natural variation. This study not only highlights the potential of high-throughput precision genome editing to resolve foundational questions in population genetics, but also to expand the exploration of empirical fitness landscapes beyond single-gene approaches toward entire biological pathways (*50*, *29*). By focusing on *natural* rather than *random* mutations, and by integrating ecological, phylogenetic, and allele-frequency information, our approach makes it possible to connect high-throughput laboratory fitness measurements to patterns of genetic variation in nature in a way that has not been possible before.

## Results

### High-throughput precision genome editing enables measurement of fitness effects for thousands of natural variants in the laboratory

To quantify the fitness effects of individual genetic variants in the Ras/PKA and TOR/Sch9 pathways we employed CRISPEY-BAR+, a modified version of a previously published high-throughput CRISPR-based precision genome editing technology. This previous method, called CRISPEY (*26*), provides a means to introduce many predefined mutations into a yeast population, one per cell, through a plasmid library of guide RNAs and corresponding donor templates for each. In this system, the donor DNA is generated *in vivo* by a bacterial retron element, which reverse-transcribes the donor RNA into single-stranded DNA that serves as a repair template for Cas9-induced double-stranded breaks. As a result of the unique biochemistry of retrons, the donor DNA is covalently linked to the RNA molecule that contains the guide RNA, ensuring its co-localization with the Cas9/gRNA complex. Critically, because the method has such high editing efficiency, it is possible to infer the frequency of the edit within a population by measuring the abundance of the corresponding guide/donor, thereby enabling pooled measurements of fitness via high-throughput sequencing of the plasmids. An improvement of the original method, CRISPEY-BAR, is based on a new vector that incorporates two consecutive retron-guide-donor cassettes, allowing the simultaneous generation of two guide/donor pairs for making two precise edits in the same cell (*29*, *30*). One guide/donor pair integrates a genomic barcode in a neutral locus, and the other introduces a precise edit of interest elsewhere in the genome. With our new method CRISPEY-BAR+ we also generate one precise edit and a unique barcode per cell and, in a pooled assay, measure the change in abundance of the edited strain through deep sequencing of the chromosomal region where barcodes are integrated (Fig. 1A). However, in contrast to CRISPEY-BAR (*29*, *30*), every guide/donor pair, and thus the variants that they introduce, is associated with hundreds of unique barcodes (Fig. S1A), providing robust internal controls to ensure changes in fitness are linked to genome editing and improving the precision of our subsequent fitness measurement (Fig. S1B). The association between barcodes and edits is established by sequencing the plasmid library (final plasmid library in Figure 1A; Methods).

**Figure 1.**
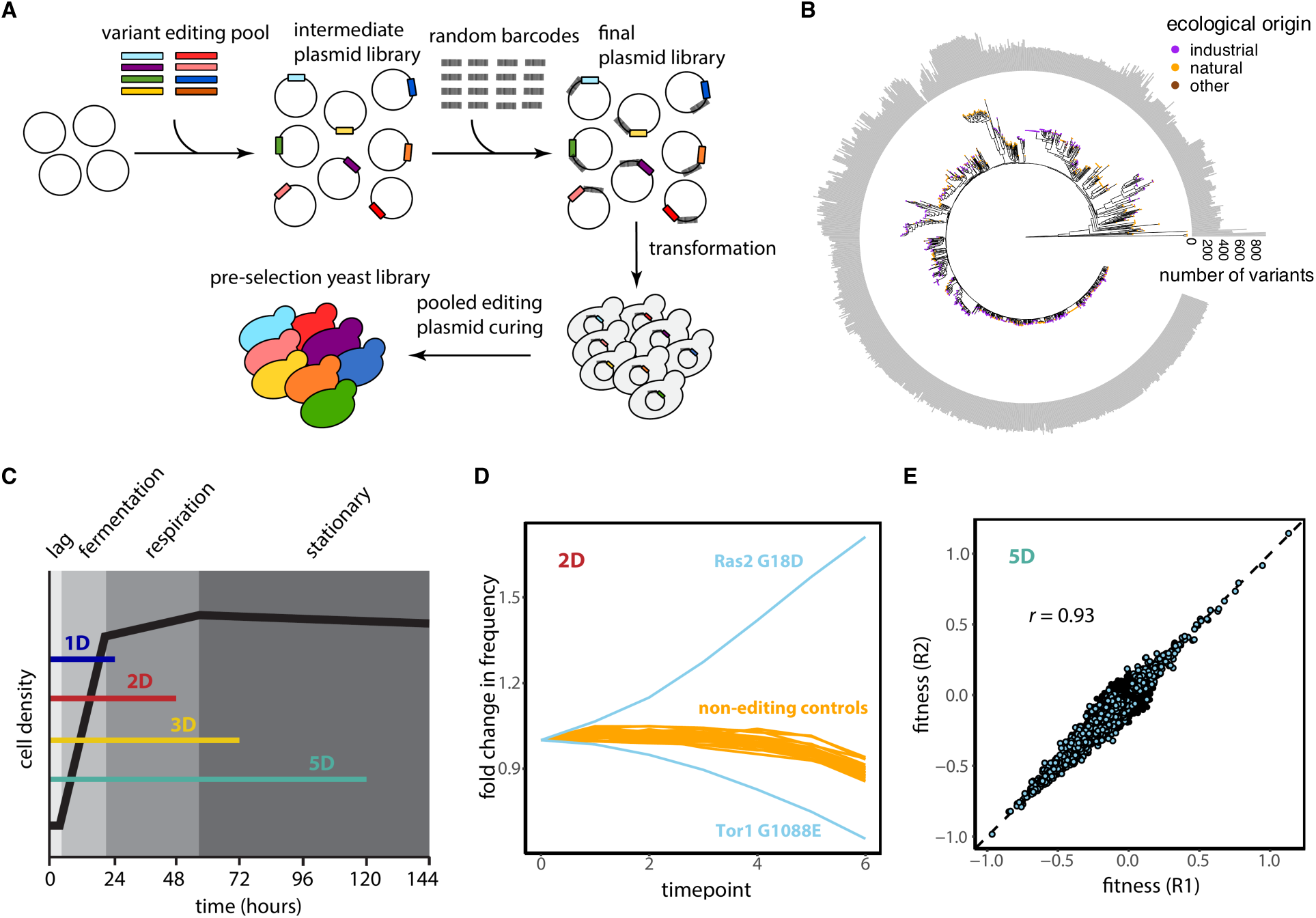
CRISPEY-BAR+ fitness screen of natural genetic variants in nutrient-sensing pathways. (A) Schematic of the CRISPEY-BAR+ approach: from cloning to pre-selection yeast population. The association between random barcodes and editing oligos is obtained by sequencing the final plasmid library. (B) Maximum likelihood tree of the 1,011 yeast strains. Individual strains are labeled according to their ecological origin, based on the provenances reported by Peter *et al.* (*20*), as industrial, natural, or other (see Methods). We report the number of variants in the Ras/PKA and TOR/Sch9 pathways for each strain. (C) Schematic of the four experimental conditions with varying time before transfer (1D: 24 h, 2D: 48 h, 3D: 72 h, 5D: 120 h) to change the amount of time spent in different phases of the growth cycle (lag, fermentation, respiration, and stationary). (D) Example of competition data over time from the 2D condition. Each blue line represents the average across three experimental replicates of the normalized counts for two missense edits: a beneficial (Ras2 G18D) and a negative (Tor1 G1088E). The beneficial variant is adjacent to the most common G12X oncogenic mutation in humans (*RAS2* is an ortholog of *KRAS, NRAS, and NRAS*) (*53*). Counts in later time points are normalized to the counts in the first time point. All the barcodes mapping to the same edit are counted. Orange lines represent the normalized counts for barcodes associated with non-editing oligos, representing a neutral fitness baseline. These orange lines go down in frequency because as the mean fitness of the population increases as a consequence of the increase in frequency of beneficial variants. (F) Estimates of relative fitness per cycle are highly reproducible across technical replicates. The scatter plot shows fitness values across replicate 1 (R1) and replicate 2 (R2) for each variant in the 5D condition. Non-neutral variants—those with significant fitness effects—are highlighted in blue, while neutral variants are shown in black. The dashed diagonal line indicates the identity line, and the Pearson correlation coefficient (*r*) quantifies replicate agreement. See Figure S2 for all replicate comparisons across all conditions.

After curating the relevant literature, we selected 18 genes from the Ras/PKA pathway and 6 from the TOR/Sch9 pathway (Supplementary Table S1), focusing on those most central to these signaling pathways. Using genome-wide polymorphisms identified from whole-genome sequences of 1,011 genetically and ecologically diverse yeast isolates (spanning wild and domesticated strains) (*20*), we identified 14,213 variants segregating in *S. cerevisiae* within these pathways, including both single-nucleotide polymorphisms (SNPs) as well as indels. Most (9,953) were in one of the 24 protein-coding regions, while the remaining 4,260 were in adjacent noncoding regions.

We were able to target 9,589 of those 14,213 variants with CRISPEY-BAR+, based on their proximity to a PAM site required by Cas9. Across edits, we observed an average editing frequency of ∼85% determined from randomly picked barcoded strains (Methods), which is in agreement with previous studies using the CRISPEY editing technology (*28*, *29*). The 9,589 variants are broadly distributed across the 1,011 strains (Fig.1B), and span a wide range of functional types: 37.9% (3,629) are nonsynonymous, 36.2% (3,473) are synonymous, 24.5% (2,349) are noncoding, and the remaining 1.4% (138) are either frameshifts, insertions/deletions in coding regions, or impacted start/stop codons.

To investigate fitness in an environment linked to the function of the Ras/PKA and TOR/Sch9 pathways, we competed the edited pool in a minimal medium with limiting glucose (1.5%, M3) over ∼25 generations in serial batch culture (*51*). We conducted growth competitions under four experimental conditions, each with three technical replicates. The experimental conditions varied in the duration the cells spent in the flask before transfer: 24 hours (1D), 48 hours (2D), 72 hours (3D), and 120 hours (5D) (Fig. 1C). As the growth cycle lengthens, cells transition from active fermentation (1D) to fermentation and respiration (2D) to prolonged stationary phase at 3D and 5D, once the limiting nutrients are depleted. Previous work with adaptive mutants in the Ras/PKA and TOR/Sch9 pathways, conducted in laboratory evolution experiments, demonstrated that these growth conditions should enable us to reveal the fitness effects of functional variation in these pathways (*42*, *46*).

During the growth competitions, we sampled cultures every ∼4.3 generations. We employed high-throughput sequencing of the barcode region in these samples to monitor the relative abundance of variants over time (Fig. 1D). We then used these frequency trajectories to estimate fitness effects of each variant. To control for the distribution of neutral fitness measurements that arise in wild-type cells from experimental noise and genetic drift, we measured variant fitness effects from each competition relative to non-editing control plasmids that do not alter the genome apart from the barcode insertion (Fig. 1D). We used a Bayesian approach to quantify the selection coefficients for each variant (*52*). This approach is based on a hierarchical model that employs all experimental replicates simultaneously for the inference of fitness (Methods). We estimated selection coefficients for 9,447 variants out of the 9,589 that we could target with CRISPEY-BAR+. The remaining 142 variants lacked sufficient barcodes in the competition data to assess their fitness. We found that the fitness effects measured by CRISPEY-BAR+ were highly reproducible across replicates, with an average Pearson’s *r* of 0.93 across conditions and pairwise replicate comparisons (*P* < 2.2 × 10^-16^; Fig. 1E and Fig. S2). To validate that observed fitness effects reflect intended edits, we analyzed control nonsense edits in cysteine or tyrosine codons of genes known to impair fitness when deleted (*26*). These nonsense edits showed significantly stronger fitness defects than synonymous edits at the same positions, supporting the accuracy and specificity of genome editing in our assay (Wilcoxon’s rank test, *P* = 6.52 × 10^-36^; Fig. S3A).

### CRISPEY-BAR+ reveals widespread fitness effects of natural variants in Ras/PKA and TOR/Sch9 signaling

To identify variants with fitness effects in our laboratory conditions, we classified a variant as significantly non-neutral if the 95% posterior credible interval for the estimated selection coefficient excluded zero (*52*). Using this criterion, we identified 3,673 (39%) variants that had significant fitness effects in at least one condition (Fig. 2A,B). Of these, 659 variants were beneficial in at least one environment (Fig. 2C,D), while the remainder were exclusively deleterious when non-neutral. Notably, 78 beneficial variants also exhibited significant adverse fitness effects in at least one other condition (Fig. 2C), revealing fitness trade-offs across different growth phases. While we refer to these variants as beneficial or deleterious here, we note that because yeast populations experience diverse and fluctuating environments in the wild, a variant beneficial under laboratory-like conditions may be mostly neutral or even deleterious in nature, depending on how frequently a lineage experiences a dissimilar type of environment. The smallest absolute fitness effect that we detected with this approach was ∼2% per transfer, providing a lower bound on the effect sizes that we can reliably resolve. The number of variants with significant fitness effects increased with longer growth cycles, peaking in the 5D condition (Fig. S3B).

**Figure 2.**
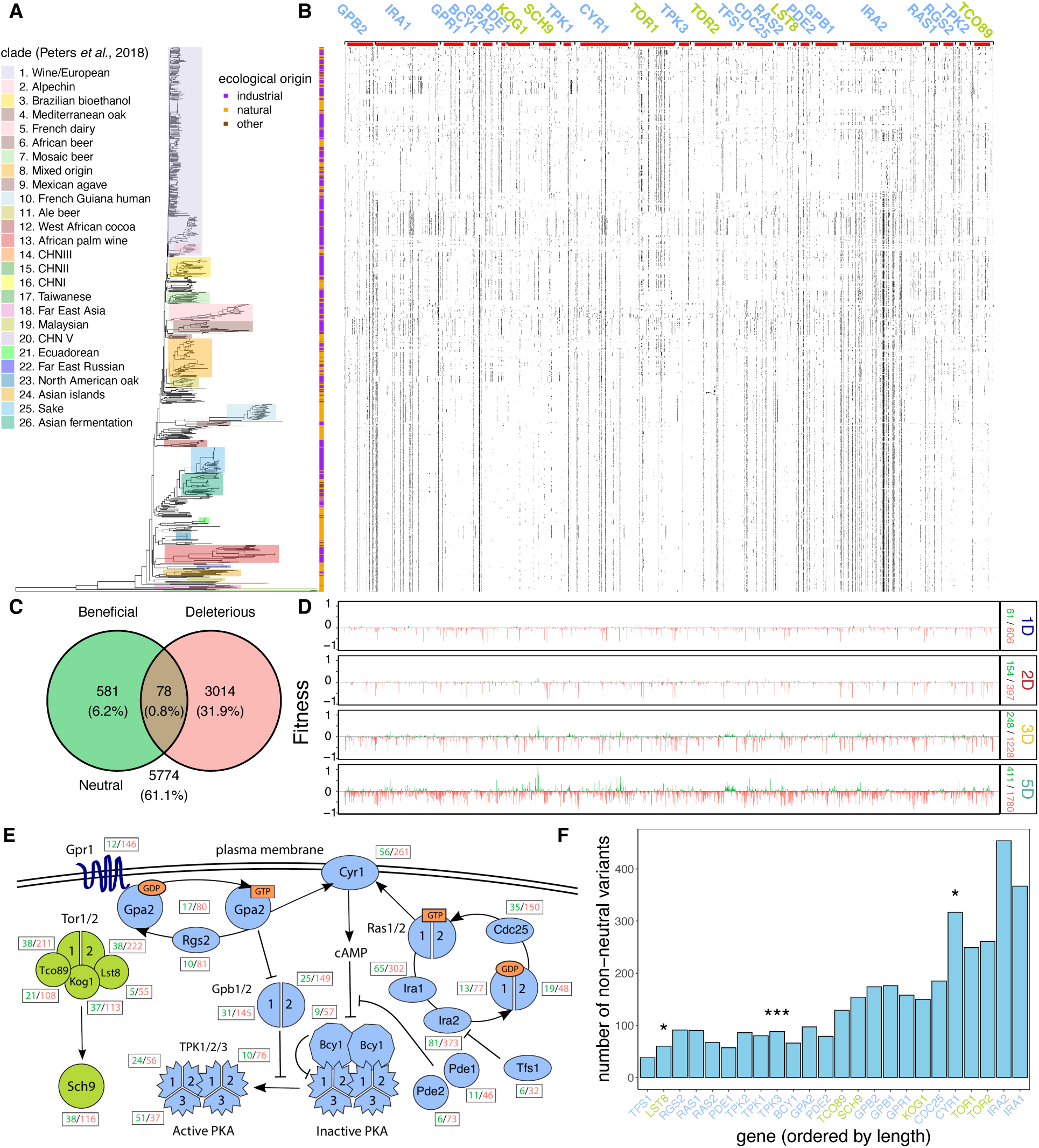
Fitness effects of segregating genetic variants in nutrient-sensing pathways revealed by CRISPEY-BAR+. (A) Maximum likelihood tree depicting phylogenetic relationship between 1,011 yeast strains (Methods). Monophyletic clades are labeled according to Peters *et al.* (*20*) and individual strains are classified according to their ecological provenance as either industrial or natural (Methods). (B) Matrix showing the presence and absence of non-neutral variants in each of the 1,011 yeast strains in the tree of panel A. Each row corresponds to a strain, and each column to a variant, ordered by genomic position. Gene annotations are indicated above the matrix, and the coding region of each gene is highlighted as a red bar. Gene names are colored by pathway: blue for Ras/PKA and green for Tor/Sch9 (following the color scheme in panel E). (C) Venn diagram showing the number of non-neutral variants that are significantly beneficial in at least one condition, significantly deleterious in at least one condition, or significantly beneficial and deleterious in different conditions (intersection). (D) Fitness of all non-neutral variants in each of the four conditions tested. Variants are colored according to panel C. The number of non-neutral variants that are beneficial (green) or deleterious (red) in each condition is listed. (E) Schematic of the signaling pathways Ras/PKA (blue) and TOR/Sch9 (green) in yeast. Positive signaling interactions are depicted with standard arrowheads and negative (inhibitory) interactions with flat-head (T-bar) arrows. The number of non-neutral variants that are beneficial (green) or deleterious (red) in each gene is listed. (F) Bar plot showing the number of non-neutral variants per gene, ordered by increasing gene length. Gene names are colored by pathway: blue for Ras/PKA and green for Tor/Sch9 (following the color scheme in panel E). Asterisks indicate genes with a statistically significant excess of variants based on a binomial test corrected for multiple testing (*FDR < 0.05; **FDR < 0.01; ***FDR < 0.001). See Figure S4 for the corresponding distribution of beneficial and deleterious variants among these pathway genes.

Non-neutral variants are widespread across both the Ras/PKA and TOR/Sch9 pathways, with no entire gene devoid of them (Fig. 2E and Fig. S5). To assess whether specific genes carry more non-neutral variants than expected given the number of assayed variants on them, we performed binomial tests, accounting for the overall background rates of fitness-altering variation within each pathway (Benjamin-Hochberg false discovery rate, FDR < 0.05). We identified three genes with a significant excess of non-neutral variants (Fig. 2F): *CYR1,* which encodes the cAMP-generating adenylate cyclase; *TPK3*, a subunit of the cAMP-dependent protein kinase A (PKA); and *LST8*, a regulatory subunit shared by both the Tor1-containing TORC1 and the Tor2-containing TORC2 complexes. When separating non-neutral variants further by beneficial or deleterious, we found that beneficial variants were particularly enriched in the terminal effector kinases of each pathway (Fig. S4A): kinase Sch9 (*SCH9*) in the TOR/Sch9 pathway and PKA (*TPK3*) in the Ras/PKA pathway. These two genes (*SCH9* and *TPK3*) alone account for approximately 12% of all beneficial variants detected (Fig. 2E). This enrichment may reflect the fact that these kinases are positioned at the downstream ends of their respective signaling cascades, where they directly control growth-related transcriptional and metabolic programs. As terminal effectors, mutations in these genes may offer a fine-tuned or conditionally advantageous means of modulating pathway output, potentially making them frequent targets of adaptation.

### Non-neutral variants tend to be nonsynonymous and are predicted to be functional

We next sought to investigate the predicted molecular impacts of the variants that show significant fitness effects in our CRISPEY-BAR+ measurements, leveraging the single-nucleotide resolution of our approach. Non-neutral variants are widely distributed across all functional categories of variants (Fig. S3C). In line with prior expectation of functionality, large fractions (∼60%; 25/42) of variants that abolish start/stop codons or create new stop codons, and ∼44% (1,612/3,629) of all non-synonymous variants are significantly non-neutral, in both cases, this is a significantly higher proportion than either synonymous (∼33%; 1,134/3,473) or noncoding (∼37%; 866/2,349) variants, which are generally less likely to impact function (Fisher’s exact test, *P* << 0.001 in all cases; Fig. S3C). Moreover, the overrepresentation of nonsynonymous variants amongst non-neutral variants increases with the number of conditions in which a variant is non-neutral (Fig. 3A, Fig. S5A, and Fig. S6A), and non-synonymous variants have on average substantially larger fitness effects than synonymous variants (Wilcoxon’s test, *P* = 1.7 × 10^-7^; Fig. S3D). Consistent with a previous study (*54*), non-neutral synonymous variants tend to decrease codon optimality specifically near the start of genes relative to neutral variants (Fig. S7). This may reflect changes to the *ribosome ramp*—a region of suboptimal codons near the 5’ end of coding sequences thought to reduce ribosome crowding and optimize translation efficiency (*54*).

**Figure 3.**
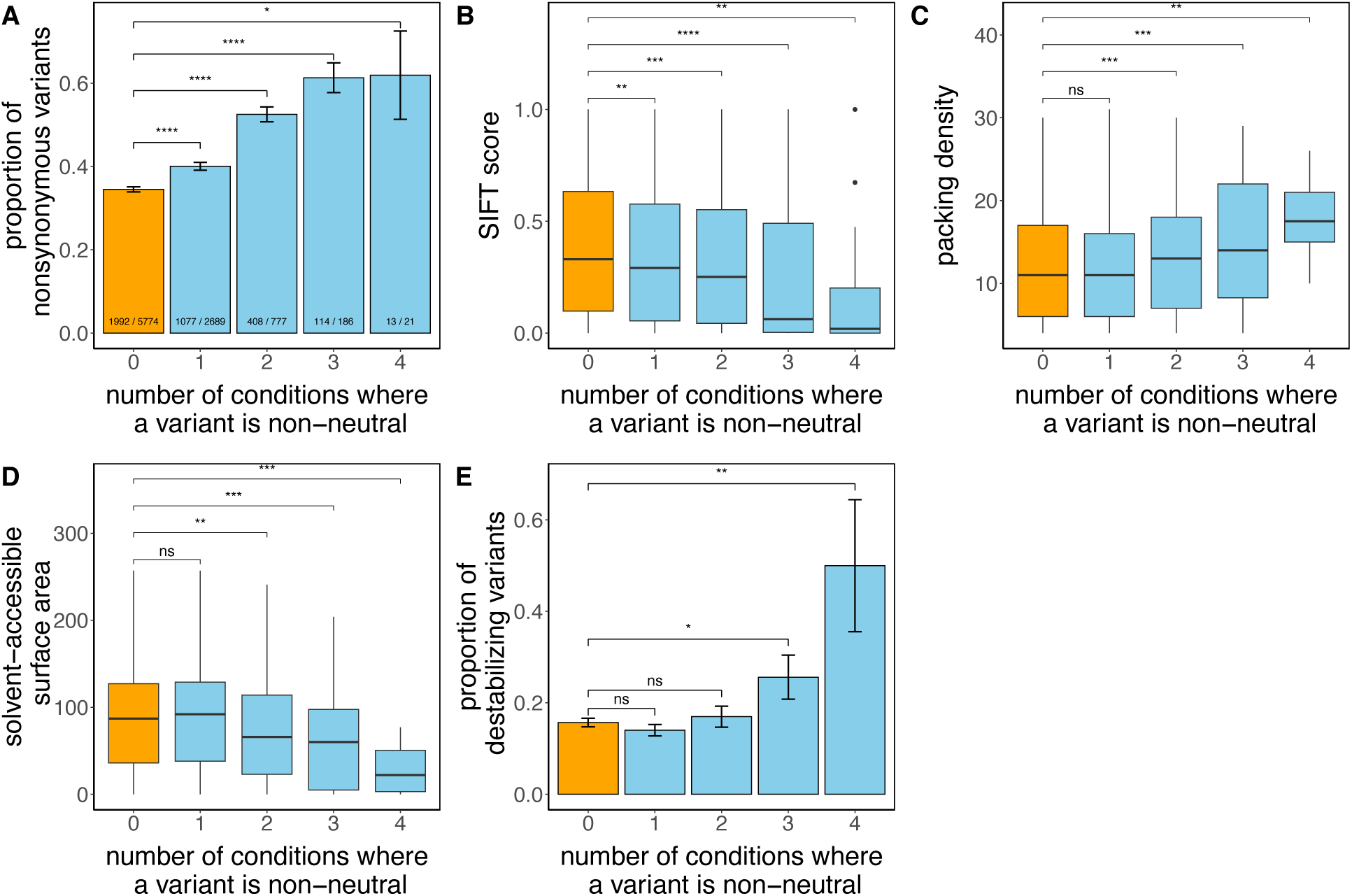
Non-neutral variants tend to be missense and are predicted to be functional. (A) Bar plot showing an enrichment of nonsynonymous variants among non-neutral variants compared to neutral variants. In this and all subsequent panels, non-neutral variants are grouped by the number of conditions in which they exhibit non-neutral fitness effects. The ratios indicate the number of nonsynonymous variants among all the variants of each class. (B) Box plot showing that non-neutral variants tend to affect more evolutionarily conserved residues, as indicated by lower SIFT scores. (C) Box plot showing that non-neutral variants tend to affect residues with higher packing density. (D) Box plot showing that non-neutral variants tend to affect residues that are less exposed to the solvent. (E) Proportion of destabilizing variants for non-neutral and neutral variants. Differences between non-neutral and neutral variants were assessed using Fisher’s exact test for panels A and E, and the Wilcoxon test for the remaining panels. Significance levels are indicated as follows: *P* ≤ 0.0001: ****, *P* ≤ 0.001: ***, *P* ≤ 0.01: **, *P* ≤ 0.05: *, not significant: ns. Box plots show the median an interquartile range; whiskers extend to 1.5 times the interquartile range.

We then proceeded to evaluate how specifically nonsynonymous non-neutral variants affect different protein properties. To do so, we related the number of conditions in which a nonsynonymous variant is non-neutral to functional predictions based on evolutionary conservation or protein structure and stability (Fig. 3B-E, Fig. S5B-E, and Fig. S6B-E). Variants that are non-neutral in multiple environments are expected to have larger, more robust molecular consequences, rather than condition-specific or weak effects.

We first found that if a missense variant occurs within a conserved protein region it is more likely to have an impact on fitness. For this analysis, we used Sorting Intolerant from Tolerant (SIFT) scores, which predict whether an amino acid substitution is likely to affect protein function based on sequence conservation and the physical properties of the amino acids involved (*55*, *56*). We find that non-neutral nonsynonymous variants tend to have lower SIFT scores than neutral nonsynonymous variants suggesting they are more likely to affect protein function (Wilcoxon’s test, *P* = 1.1 × 10^-7^; Fig. 3B, Fig. S5B, and Fig. S6B). Variants with a SIFT score below 0.05 are considered to have particularly strong functional impacts (*55*). There is an enrichment of such variants among non-neutral variants in our assays (Fisher’s test, *P* = 1.6 × 10^-8^).

We next examined three key protein properties: packing density, solvent accessibility, and protein stability. Packing density reflects how closely a residue is located to others within the protein’s tertiary structure, essentially measuring the degree of contact a residue has with other residues. Solvent accessibility represents the surface area of a residue that is exposed to water. Substituting residues in densely packed regions, as well as those with lower solvent exposure, is likely to induce considerable changes to the protein’s structure and are therefore more evolutionarily constrained (*57*). Indeed, we find that nonsynonymous variants that are non-neutral in more environments tend to affect residues with higher packing density (Fig. 3C, Fig. S5C, and Fig. S6C) and those that are less solvent-exposed (Fig. 3C, Fig. S5C, and Fig. S6C). As a control, we confirmed that there were no significant differences in packing density or solvent accessibility comparing codons containing non-neutral relative to those containing neutral synonymous variants (Fig. S8). Because nonsynonymous variants in these structurally constrained sites often destabilize the folded protein, we next asked whether non-neutral variants are more likely to impair stability. Using FoldX to estimate changes in folding free energy (ΔΔG) (*58*), we found an enrichment of strongly destabilizing variants (ΔΔG > 2) among those that are non-neutral in multiple environments (Fig. 3E, Fig. S5E, and Fig. S6E).

In summary, our findings indicate that non-neutral variants are more likely to be missense mutations, affecting conserved, densely packed, and buried residues within a protein, and often resulting in destabilization. These results provide a strong biochemical basis for the distinction between neutral and non-neutral variants, further validating the findings of our high-throughput fitness measurements.

### A substantial fraction of non-neutral variants shows high allele frequency

Having established that non-neutral variants in our CRISPEY-BAR+ assay have larger molecular impacts than neutral variants, we next set out to explore their allele distribution across 1,011 natural isolates of *S. cerevisiae* spanning the global ecological diversity of this species.

Nearly neutral theory predicts that common and older variants should have only weak effects on fitness, while rare and young variants can be deleterious and maintained in a mutation– selection balance. In contrast to this expectation, we often observe non-neutral variants across all frequency classes: of the 3,673 variants that are non-neutral in at least one condition, 887 (24.1%) are common (present in more than 10 strains). Of these 887 common non-neutral variants, 122 (13%) have a maximum absolute fitness across all four conditions that is larger than twice the average (∼33% per transfer) (Fig. 4A). Furthermore, we detect no enrichment of rare variants among large-effect non-neutrals (Fisher’s exact test, odds ratio = 0.94, *P* = 0.617) and there is only a weak negative relationship between maximum absolute fitness and the number of strains in which a variant is found (Spearman’s ρ = −0.052, *P* = 1.6 × 10^-3^).

**Figure 4.**
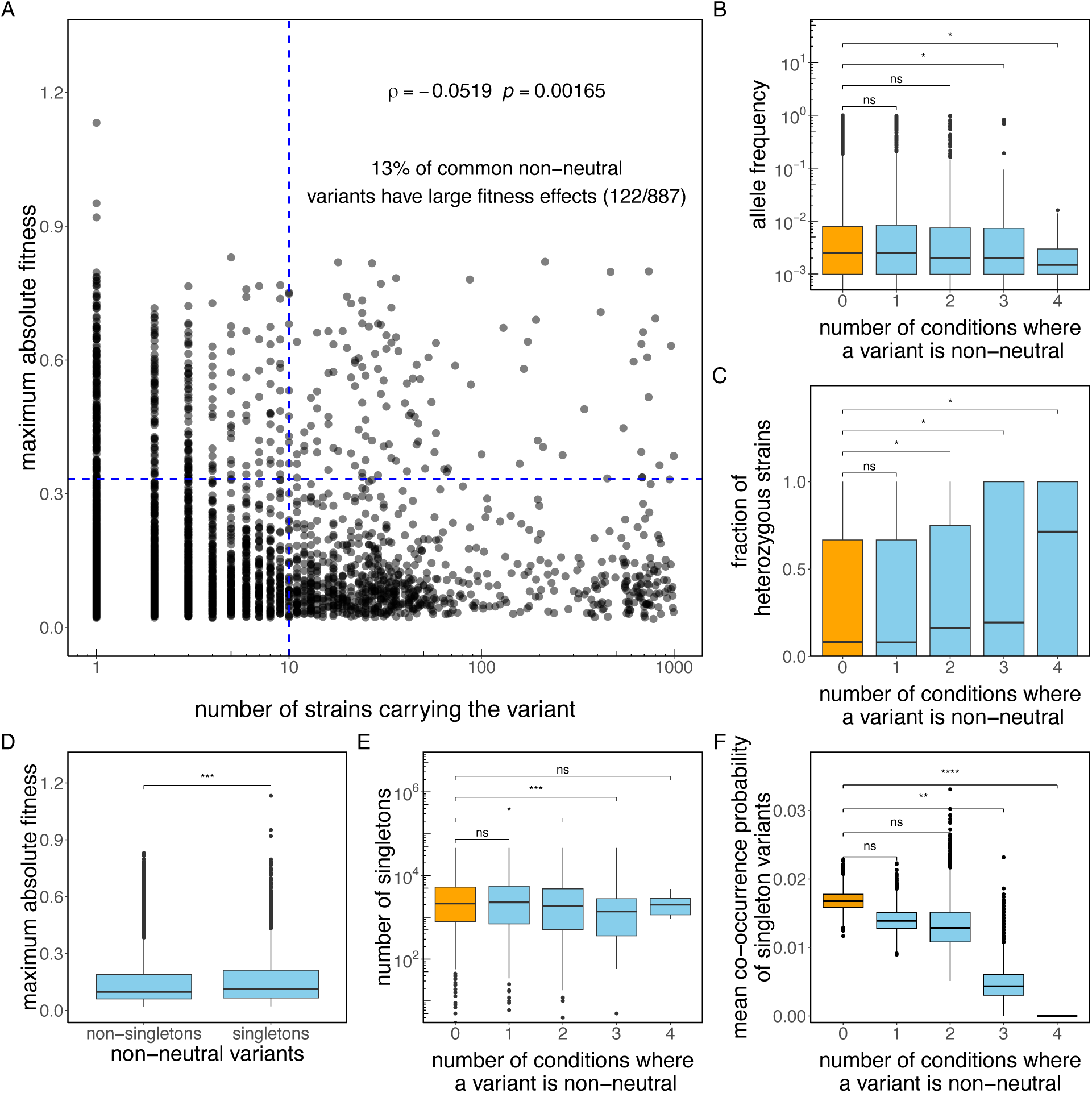
A substantial fraction of non-neutral variants shows high frequency, while low-frequency variants show evidence of purifying selection. (A) Scatter plot displaying the maximum absolute fitness values across the four conditions for non-neutral variants, plotted against the number of strains in the 1,011-yeast strain collection carrying the variant. The dashed line indicates a fitness threshold set at twice the average value (0.33). (B) Box plot showing that non-neutral variants tend to have lower allele frequency. (C) Box plot showing that the fraction of strains where the variant is in heterozygosis is higher for non-neutral variants. (D) Box plot showing that the experimental fitness impact of rare variants is greater than for less rare variants. Rare variants are defined as those present in a single strain (singletons). (E) Box plot showing that singleton non-neutral variants tend to be younger. (F) Box plot showing the co-occurrence probability for neutral and non-neutral singleton variants. Non-neutral variants are grouped by the number of conditions in which they exhibit non-neutral fitness effects. The data shown represent the mean co-occurrence for 10,000 bootstrap samples (Methods). Box plots show the median and upper and lower quartiles; whiskers show 1.5 times the interquartile range.

We observed that non-neutral variants with effects across multiple conditions have slightly lower allele frequency (Spearman’s ρ = −0.039, *P* = 0.0168; Fig. 4B and Fig. S9A). These multi-condition non-neutral variants are also somewhat more likely to be found in the heterozygous state (Spearman’s ρ = 0.052, *P* = 1.6 × 10^-3^; Fig. 4C and Fig. S9B), even controlling for their lower allele frequency (partial Spearman’s ρ =0.049, *P* = 3.2 × 10^-3^). This is consistent with the expectation that deleterious variants are more likely to be recessive and have their effects masked in the largely diploid *S. cerevisiae* (*20*).

It is possible that the weakness of the statistical relationship between allele frequency and fitness effect is due to the fact that even low-frequency variants in the sample of 1,011 genomes are already sufficiently common to be largely neutral (*59*). Indeed, the class of non-neutral variants present in only a single strain (singletons) tend to have slightly larger impacts on fitness than non-singletons (Wilcoxon’s test, *P* = 2.8 × 10^-4^; Fig. 4D and Fig. S9C). Furthermore, we can approximate the age of singleton variants by the length of the terminal branch associated with the single strain in which they are found, estimated here as the total number of singleton variants present in the strain (*59*), and test for the association between the age of a singleton and their fitness impacts in our measurements. Although we do not find a significant correlation between maximum absolute fitness and the estimated age of a non-neutral singleton variant (Spearman’s ρ = −0.022, *P* = 0.450), we observe that compared to neutral singletons, singletons having non-neutral effects in multiple conditions tend to be younger (Spearman’s ρ = −0.1, *P* = 6.6 × 10^-4^; Fig. 4E and Fig. S9D). For example, singletons that are non-neutral in at least two conditions are disproportionately found in less divergent strains which on average harbor 1,688 fewer strain-specific variants (6,015 vs. 7,703; Wilcoxon’s test, *P* = 2.7 × 10^-4^).

Additionally, non-neutral singletons tend to co-occur within the same strain less frequently than neutral singletons as the number of non-neutral conditions in which they are non-neutral increases (*P* < 1 × 10^-4^, bootstrap test for Spearman’s ρ < 0, Fig. 4F and Fig. S9E). For example, singletons that are non-neutral in at least two conditions co-occur 1.65-fold less often than neutral singletons (bootstrap test for difference in mean co-occurrence > 0, *P* < 1 × 10^-4^). This is consistent with purifying selection, which continuously purges deleterious variants before multiple can accumulate within a single genome.

These analyses suggest the existence of a subclass of rare functional variants maintained at low frequency in the population under mutation-selection balance, consistent with classical, nearly neutral theory. However, despite these detectable albeit weak signatures of purifying selection, we also find clear evidence that many non-neutral variants persist at high frequencies. Thus, while rare variants are modestly enriched for stronger fitness effects in laboratory conditions, a substantial fraction of common polymorphisms is functionally impactful, and at times strongly so.

### Beneficial variants show evidence of local adaptation

The discovery that common variants can have very large fitness effects in laboratory conditions is consistent with balance theory, in which these variants are driven to high frequencies by local adaptation. However, the fitness effect of variants in experimental conditions may not reflect the fitness effect in more complex natural conditions. To provide evidence for local adaptation, it is essential to associate fitness effects in our assay with ecological variables. We therefore next set out to explore whether non-neutral variants are more commonly found in specific clades or associated with specific ecological niches.

*S. cerevisiae* is the dominant species in the fermentation of various beverages and foods and has been domesticated on multiple independent occasions (*60*, *61*). The process of domestication has had a large impact on the stress tolerance and the fermentative growth capacity of this species, which are traits regulated by the core nutrient-sensing pathways we study here (*62–64*). We therefore hypothesized that natural genetic variation in these pathways may have contributed to adaptation in industrial environments.

We first focused on the 2,851 variants we assayed that are found exclusively in strains from industrial environments (industrial variants), and compared them to the 3,250 variants solely found in natural strains (Methods). As expected, industrial variants were modestly enriched for non-neutral effects (Fisher’s exact test, odds ratio = 1.125, *P* = 0.025). This enrichment became substantially stronger when focusing on variants that are beneficial in at least one condition (odds ratio = 1.56, *P* = 9 × 10^-6^). It disappeared when examining non-neutral variants that are exclusively deleterious (Fisher’s exact test, odds ratio = 0.99, *P* = 0.91) (Fig. 5A). These findings suggest that industrial strains are not merely enriched for functional variants in general, but specifically for variants which are beneficial under the laboratory conditions we tested which have previously been shown to select for Ras/PKA and TOR/Sch9 variants strongly impacting fermentative performance, respiration and starvation tolerance (*46*).

**Figure 5.**
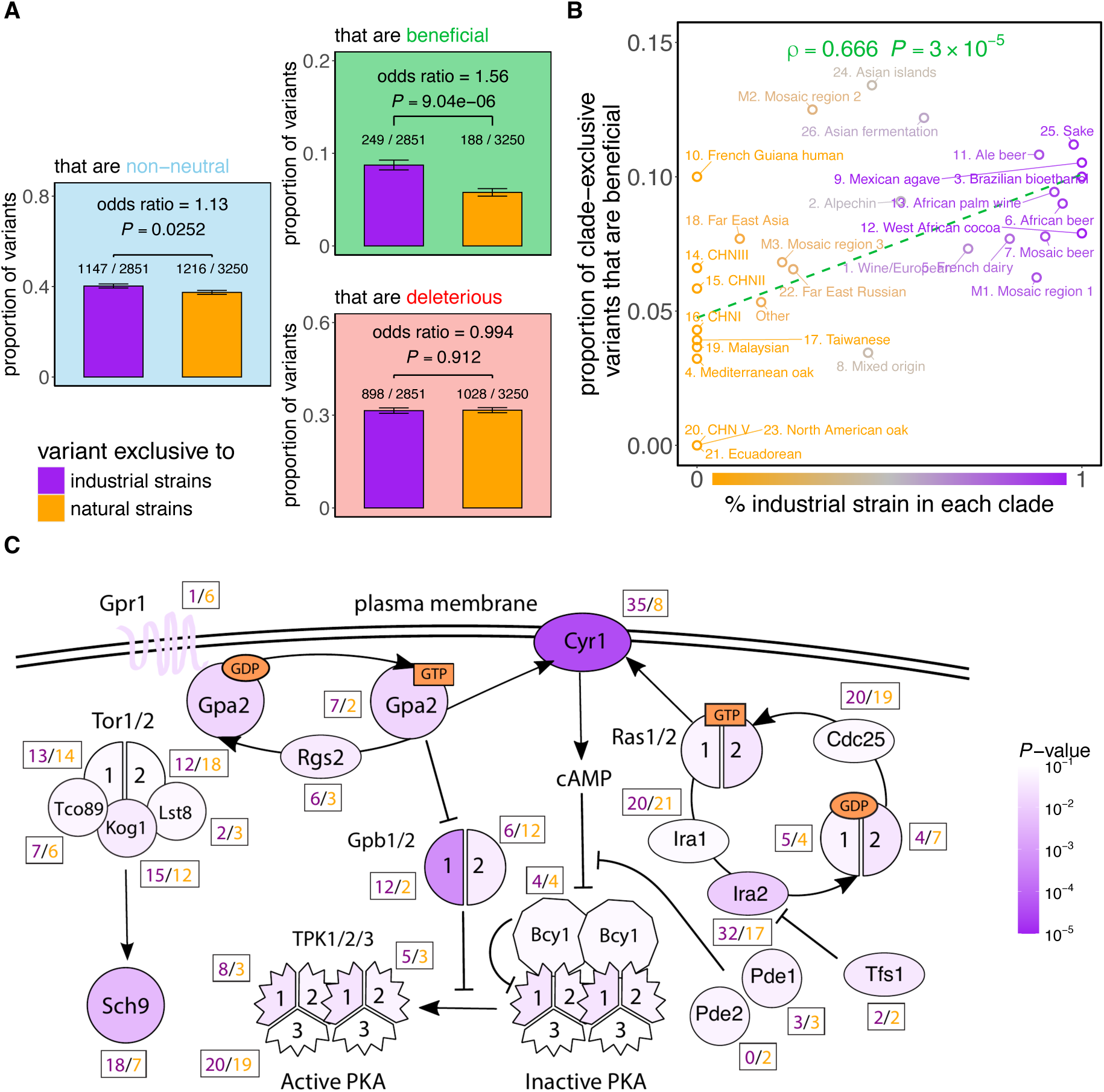
Beneficial variants show evidence of local adaptation. We classify variants according to whether they are exclusively observed in yeast strains from natural origins (orange) or industrial origins (purple) (Methods). (A) Bar plots comparing the proportion of variants in each group that are non-neutral (blue), beneficial (green) or deleterious (red). Each of the 26 clades shown in Figure 2A has several clade-exclusive variants, and many contain a mixture of industrial and natural strains. (B) Scatter plot displaying the proportion of clade-exclusive variants that are beneficial against the proportion of strains in each clade that are from an industrial origin. (C) Schematic of the signaling pathways Ras/PKA and TOR/Sch9 in yeast. The number of beneficial variants that are exclusively industrial (purple) or natural (orange) for each gene is listed. Genes are colored by the odds-ratio *P*-value (FDR-corrected) of the enrichment for exclusively industrial variants (Fig. S11B).

Disentangling the role of ecology from shared ancestry is challenging because of the phylogenetic structure of the 1,011-strain panel (Fig. 2A): closely related strains often share both habitats and variants. To address this, we examined variants restricted to individual clades and related their beneficial effects to the proportion of industrial strains within each clade.

Variants in clades with more industrial strains were significantly more likely to be beneficial (Spearman’s ρ = 0.666, *P* = 3 × 10^-5^) (Fig. 5B). To test whether this association held within clades, we leveraged the fact that many clades contain both industrial and natural strains. We used a logistic mixed-effects model to quantify the effect of a strain having an industrial origin on the likelihood that a variant is beneficial with the clade identity as a random effect (Methods).

Within clades, variants found exclusively in industrial strains are significantly more likely to be beneficial than those found exclusively in natural strains (effect size = 0.43, *P* = 8.8 × 10^-4^). Our model shows that the observed enrichment of beneficial variants persists even after controlling for shared evolutionary history, suggesting local adaptation within industrial environments.

Furthermore, the enrichment of beneficial variants in industrial strains is significantly larger than in null models, which control for systematic variation in allele frequency and SIFT scores associated with industrial variants along the phylogeny (Methods; S10A-B and Fig. S11A-F).

One possible confounder is that domestication may have relaxed purifying selection via population bottlenecking, allowing deleterious variants to rise in frequency in industrial strains irrespective of whether they are in the Ras/PKA or TOR/Sch9 pathways (*65*). Indeed, consistent with this genome-wide bottlenecking and relaxation of purifying selection, nonsynonymous variants exclusive to industrial strains tend to have lower allele frequency and SIFT scores than variants exclusive to natural strains, regardless of whether they are in the Ras/PKA or TOR/Sch9 pathways (Fig. S10C-D). To account for this, we generated 10,000 matched control sets of variants outside the signaling pathways we studied, with equivalent allele frequency and SIFT score distributions to the non-synonymous variants in our Ras/PKA and TOR/Sch9 CRISPEY-BAR+ screen (Fig. S11G-I). For each control set, we quantified the enrichment of beneficial variants as assessed by the odds ratio (Fig. 5A), clade-level correlation (Fig. 5B), and the mixed-effects model (Methods). These controls failed to reproduce the observed enrichment of beneficial Ras/PKA and TOR/Sch9 CRISPEY-BAR+ variant effects, suggesting that the signal we detect is not simply due to bottlenecking but instead reflects Ras/PKA- and TOR/Sch9-specific local adaptation to industrial environments.

Finally, we set out to identify the nature of the beneficial variants that are enriched in industrial strains. We first asked whether these beneficial and exclusively industrial variants would be confined to one of the two major pathways or to a specific subset of genes. We found that the enrichment holds for variants in both the Ras/PKA and TOR/Sch9 pathways when analyzed independently (Fig. S12A). These variants are found across a broad range of genes in the pathways, affecting 23 of the 24 genes we have studied (Fig. 5C and Fig. S12B). Three genes show a significant enrichment of such industrial and beneficial variants as assessed by the odds ratio, clade-level correlation, and the mixed effect model (Fig. S12B): the adenylate cyclase *CYR1*, the effector kinase *SCH9*, and *GPB1*, a negative regulator of cAMP/PKA signaling (but not its paralog *GPB2*). Of these, only *SCH9* is significantly enriched for beneficial variants more generally (Fig. S4B), indicating that the enrichment in industrial strains reflects selection for specific alleles that are advantageous in industrial environments rather than simply increased statistical power when genes carry more variants that are beneficial in our pooled competition assays.

These results leave open the question as to whether these signatures of local adaptation are unique to the Ras/PKA and TOR/Sch9 pathways or occur more broadly across the genome. Across these two pathways, 3.67% of nonsynonymous variants are present in more than 10 strains and had a SIFT score below 0.05—a proportion significantly higher than the genome-wide average of 2.6% (two-proportion Z-test, *P* = 6.45 × 10^-6^; Fig. S13A). This enrichment was robust to the thresholds used to define common and functional variants, supporting the idea that Ras/PKA and TOR/Sch9 are potential genomic hotspots of local adaptation (Fig. S13B).

Nevertheless, many other genes outside these pathways also showed strong enrichment for common variants with predicted impact on molecular function (Fig. S13C).

We also find that functional variants (SIFT score < 0.05) found exclusively in industrial strains are enriched in the Ras/PKA and Tor/Sch9 pathways relative to the rest of the genome, whereas this is not true for exclusively natural variants, consistent with the idea that these signaling pathways underpin local adaptation to industrial environments (Fig. S13D). After correcting for multiple testing, we identified 26 genes with significant enrichment for functional, industrial variants (Fig. S13E). Notably, several of these genes have been previously implicated in industrial adaptation (*49*, *66*). Gene-set enrichment analysis suggests that these genes are disproportionately involved in carbohydrate metabolism—a key trait likely advantageous in nutrient-rich, stress-prone industrial fermentations (Fig. S13E). Importantly, this genome-wide enrichment analysis lacks the statistical power to recover the Ras/PKA and TOR/Sch9 genes enriched for beneficial and exclusively industrial variants, suggesting that many other genes may be involved in local adaptation to industrial fermentation environments. This demonstrates why identifying functional variants with CRISPEY-BAR+ and then testing their ecological enrichment is essential for detecting genomic hotspots of local adaptation that would otherwise be missed if we rely solely on predicted molecular phenotypes.

## Discussion

This study provides a comprehensive interrogation of the fitness consequences of natural genetic variation within the Ras/PKA and TOR/Sch9 pathways in *S. cerevisiae*, which are ancient and key regulators of cell proliferation across all eukaryotes. Leveraging our high-throughput CRISPEY-BAR+ approach, we precisely measured the fitness effects for over 9,000 natural variants across multiple conditions, providing insights into the evolutionary forces shaping these core pathways. The scale and precision of this experimental undertaking allowed us to tackle a central and longstanding question in population genetics: the fitness consequences of common variants in natural populations.

Some of our findings do agree with the expectations of nearly neutral theory. Namely, some of the variants that exhibit fitness effects in our experiments show signs of negative selection, as they tend to be rarer and younger, and more often found in heterozygous states, which allows for the buffering of their detrimental effects on fitness, and tend not to co-occur with each other (Fig. 4). These observations align with the idea that standing variation in these highly conserved pathways is under purifying selection, preventing the accumulation of deleterious mutations and retaining those that occur at low frequencies (*67*).

The majority of natural variants are neutral in our experimental conditions (61%). While there may be false negatives among these variants, the high overall editing rate in our screen makes it likely that most of them are indeed neutral within the limits of our detection power (∼2% per growth cycle). This result also follows the expectation of nearly neutral theory, particularly for core eukaryotic pathways that are highly conserved from yeast to humans and are therefore expected to evolve under high constraint (*68*). Nonetheless, it is remarkable that even within these highly conserved pathways, nearly 40% of variants exhibit detectable and often substantial functional consequences in at least one of the few environments we examined (Fig. 2).

Furthermore, we observed a substantial set of common (present in more than 10 strains and often at a much higher frequency) non-neutral variants with high fitness in at least one of the environments we explored (Fig. 2 and Fig. 4A). This does not naturally follow the nearly neutral expectations and shows that the high frequency of a polymorphism cannot be used to infer that it has a very weak functional impact, despite common practice (*69–72*). Moreover, we find that those variants that have beneficial fitness effects in at least one of the conditions we used, are enriched in strains associated with human-made environments, suggesting they may have contributed to local adaptation to specialized ecological niches (Fig. 5). The process of domestication has significantly influenced the stress tolerance and fermentative growth capacity of *S. cerevisiae*, which are precisely the traits affected by the Ras/PKA and TOR/Sch9 pathways (*73*). This aligns well with the idea that local positive selection has affected variation in these nutrient-sensing pathways. Another reason for the strong signal of local adaptation we uncover may be that the relatively stable and controlled environments experienced by domesticated yeast strains offer less ecological complexity compared to wild environments, where mutations that lock signaling pathways into specific functional configurations could be highly disadvantageous across a broader range of experienced conditions (*60*).

Although laboratory conditions inevitably simplify the environments experienced by yeast in nature, some conditions can capture key components of natural environments—such as nutrient-rich, fast-growth conditions (*74*). It is therefore not unreasonable to expect some correlation between fitness measured in the laboratory and fitness in the wild. However, because natural environments fluctuate, the extent to which a variant’s experimental fitness reflects its effect in nature will depend on how frequently each lineage encounters laboratory-like conditions (*75*). This helps explain why beneficial variants are enriched in industrial strains: these strains likely experience conditions that are, on average, more similar to our experimental settings (*76*). Moreover, because adaptation can occur rapidly, there are likely ecological “pockets” where laboratory conditions provide a meaningful proxy for natural selection in the wild. If evolution were much slower, fitness effects would be averaged across many environments, and capturing them would require far more complex experimental designs.

The Ras/PKA and TOR/Sch9 pathways are also key targets of adaptation in laboratory experiments (*39*, *42*, *46*), suggesting they may have unique characteristics not shared by other signaling pathways that makes them particularly prone to accumulating local beneficial mutations. However, the signatures of domestication and local adaptation that we unveil in these two signaling pathways are unlikely to be confined to them. Indeed, we find numerous other genes with strong enrichments for exclusively industrial variants predicted to be functional (Fig. S13). Many of these are involved in carbohydrate metabolism, which is a function likely advantageous in nutrient-rich, stress-prone industrial fermentations. Notably, several of these genes have been previously implicated in adaptation to industrial environments (*49*, *66*), underscoring that domestication in *S. cerevisiae* is polygenic and shaped by multiple adaptive routes. Therefore, applying our approach to additional pathways could reveal whether these systems influence adaptation in distinct ways.

Our results overall agree more strongly with balance theory than with nearly neutral theory. While most variants in these pathways are neutral in our lab environments or subject to purifying selection, we find a substantial fraction of common variation under local positive and overall balancing selection (*77*). With the advent of methods for fitness mapping *en masse* at single-nucleotide resolution, we can begin to disentangle and quantify, for the first time, the relative contributions of neutral evolution and balancing selection in shaping standing genetic variation.

Despite the strength of our findings, this study has several limitations. First, we conducted our fitness assays in a haploid background, while many yeast strains are naturally diploid. Fitness effects in diploid strains could differ for variants with recessive or partially dominant effects (*78*). Second, we focused on four environmental conditions, and exploring a broader range of conditions could reveal additional insights into the pleiotropic effects of these variants (*29*). Third, fitness effects smaller than our lower limit of detection (∼2% per transfer) could still be highly impactful in the large populations often found in both natural and industrial environments. Finally, by measuring the fitness of individual variants in a single genetic background, we could not investigate the potential role of epistasis in modulating variant fitness across different genetic contexts and environments (*79*, *30*, *80*, *81*). Taken together, these four limitations suggest that the true number of variants with functional consequences may in fact be greater than what we estimate here.

Future studies using our method could address these limitations by measuring fitness in a wider range of strains, including diploid strains, under a broader range of environmental conditions, such as more complex media that mimic natural ecological conditions. However, despite all the complexities that we did not explore in this study, one of its more surprising outcomes is the strong correlation between fitness effects measured in the laboratory and ecological adaptations observed in nature. Our work demonstrates that laboratory-based fitness assays can be relevant to real-world evolutionary outcomes. Looking ahead, integrating massively parallel fitness measurements with modern machine learning approaches offers exciting opportunities to build more accurate, mechanistic, and predictive models of how genomic information, at nucleotide resolution, gives rise to phenotypes and fitness (*82*).

In conclusion, this study bridges the gap between population genetics and functional genomics by providing direct measurements of fitness effects for natural variants across entire pathways. We constructed a comprehensive pathway-wide, nucleotide-level fitness landscape of natural variation, demonstrating how massively parallel fitness mapping methods can be used to address long-standing questions in evolutionary biology by leveraging ecological, phylogenetic, and allele-frequency information. Our findings are more consistent with predictions of balance theory than with nearly neutral theory, highlighting how natural selection can actively maintain functional genetic diversity in key signaling pathways. Future studies that expand this approach to additional pathways, environments, and genetic backgrounds will further our understanding of the genotype-fitness map, providing a more nuanced view of the evolutionary dynamics that shape natural populations.

## Acknowledgments

This work was supported by the Swiss National Science Foundation (P2ZHP3_174735 to J.A.-R.), the European Molecular Biology Organization (ALTF 724-2018 to J.A.-R.), the CZ Biohub (to D.A.P.), the National Institutes of Health (5R35GM118165-07 to D.A.P.; GM156526 to H.B.F.; DP2-GM119140, RF1-AG057334, R01-AG06341801, and R01-HG012366 to D.F.J.), the National Science Foundation (NSF-MCB116762 to D.F.J.), a Searle Scholar Award (14-SSP-210 to D.F.J.), a Kimmel Scholar Award (SKF-15-154 to D.F.J.), a Vallee Scholar Award (to D.F.J.), and a Discovery Innovation Award from Stanford University (to D.F.J.). We are grateful to Roy Ang, Stefan Oliver Bassler, Shreyas Gopalakrishnan, Chris Jakobson, Bernard Kim, Grant Kinsler, Alexandra Khristich, and Alan Su, and all other members of the Petrov Lab and Jarosz Lab for discussions during experimental design, experimental work, and data analysis.

## Author contributions

J.A.-R. and D.A.P. conceived the project. J.A.-R. conducted experiments, collected data, and performed analysis. D.A.P. and D.F.J. provided supervision and advised on experimental design and data analyses. S.A.C and H.B.F. provided extensive feedback on experimental design and helped with experimental troubleshooting. J.A.-R., J.C.C.V. and D.A.P. interpreted results. J.C.C.V. conducted phylogenetic analyses and analyses related to evidence for local adaptation, prepared figures, and wrote the results and methods sections for those analyses. J.A.-R., M.R.-M. and O.M.G. performed fitness estimation. M.R.-M. wrote the section about fitness estimation on the Methods. J.A.-R., D.F.J., H.B.F., and D.A.P. acquired funding. J.A.-R. wrote the original draft of the paper and prepared figures. J.A.-R., J.C.C.V., and D.A.P. reviewed and edited the paper with input from M.R.-M., O.M.G., D.F.J., and H.B.F.

## Data availability

All sequence data have been deposited to SRA under NIH BioProject number (PRJNA1238315).

## Code availability

This repository contains the data and code necessary to reproduce the figures in this paper: https://github.com/jaguilarrod/signaling_pathways/

## Methods

### Media

We performed competition experiments in M3 media (*51*). This medium is glucose-limited (1.5% glucose), meaning that the cells exhaust their glucose supply before any other nutrient. This medium has been widely used in experimental evolution before (*83*, *42*, *46*, *84–88*). We supplemented M3 with uracil (20 mg/L), L-leucine (100 mg/L), and L-lysine HCl (30 mg/L) to account for the auxotrophies of ZRS111 (M3+uracil+lysine+leucine). We used SD-histidine-uracil 2% raffinose media for pre-culturing before induction: 6.7 g/L Yeast Nitrogen Base (YNB) (RPI), 1.92 g/L dropout mix synthetic minus histidine, uracil, adenine rich without YNB (US Biological), and 20 g/L raffinose (Sigma). We used SD-histidine-uracil 2% galactose media for induction, consisting of 6.7 g/L YNB, 1.92 g/L dropout mix synthetic minus histidine, uracil, and adenine rich without YNB, and 20 g/L galactose (Sigma). We also used SC media: 1.7 g/L YNB; 5 g/L ammonium sulfate (ACROS Organics), 1.9 g/L dropout synthetic mix complete without nitrogen base (US Biological), and 20 g/L glucose (Sigma).

### Variant selection and editing library design

We selected natural genetic variants from the 1,011 yeast genomes project (*20*). We selected 24 genes: 18 from the Ras/PKA pathway and 6 from the TOR/Sch9 pathway (Supplementary Table S1). For each gene, we selected variants located within the coding region as well as those within 500 base pairs upstream and downstream of the coding region, therefore including not only coding variants but also noncoding variants that could potentially impact regulation and gene expression. We designed CRISPEY oligos to edit these variants in the ZRS111 strain, which contains the S288c reference alleles. The library includes 19,906 guide/donor oligonucleotides targeting 11,058 genetic differences. We designed the guides and donor sequences in the oligos as described previously (*26*). We also included 50 pairs of oligos targeting essential genes. In each pair, one oligo introduces a nonsense mutation to a cysteine or a tyrosine codon, while the other oligo introduces a synonymous mutation in the same codon. Lastly, we designed 30 guides and donors with random sequences, which are therefore non-editing (Fig. 1D).

### Library cloning

We ordered an oligonucleotide library from Twist Biosciences. We amplified the oligonucleotide library using primers #310 and #313 with Q5 hot-start DNA polymerase (New England Biolabs). All the primer sequences used in this study are listed in Chen *et al.* (*29*). This PCR reaction produced amplicons with flanking 20 bp sequences homologous to plasmid pSAC119. We cloned the PCR-amplified double-stranded DNA into NotI-digested pSAC119 using NEBuilder HiFi DNA Assembly Cloning Kit (New England Biolabs). We performed the Gibson assembly with a molar ratio of vector to insert of 1:5.

We transformed the assembled plasmid libraries via electroporation into Endura electrocompetent cells (Lucigen). We performed five electroporations, each with 25 μL of Endura cells and 80 ng of library plasmid DNA. We conducted the electroporations in 0.1cm-gap cuvettes using a GenePulser (Bio-Rad). We allowed the transformed bacteria to recover in recovery media (supplied with Endura competent cells) for 1 hour and then plated them on LB/Carbenicillin plates for incubation at 37℃ overnight. We also plated dilutions to estimate the transformation efficiency (cfu/transformation).

We extracted plasmids using the EZNA Plasmid Maxi Kit (Omega Biotek). We eluted the plasmids in 3mL of kit-provided elution buffer. We subsequently digested the samples with NotI and CIP (New England Biolabs) to linearize the empty vectors. We then ethanol-precipitated the plasmids to concentrate the libraries at approximately 0.8 μg/μL.

The base yeast strain ZRS111 has been described previously (*26*). We performed the transformation into yeast by electroporating the plasmid library into the strain ZRS111. We prepared electrocompetent yeast by modifying previously described methods (*89*). We selected the yeast transformations by plating on SD-HIS-URA 2% agar plates. We scraped the yeast transformations off the selective plates using cell spreaders, resuspended them in SD-HIS-URA and froze them in 15% glycerol stocks at −80℃ for subsequent experiments. We generated 19 million transformants, with an average representation of 963 cells per guide/donor oligonucleotide.

### Generation of the oligo-barcode map

We amplified 120 ng of the maxiprep of the final plasmid library in a 400 μL Q5 polymerase (New England Biolabs) reaction with 1 μM forward primer #73 and 1 μM reverse primer equimolar mix of primers #327-#334. We performed the PCR reaction with 1 M betaine and an initial denaturation of 98℃ for 2 min, followed by 16 cycles of 98℃ for 10 s, 65℃ for 20 s, and 72℃ for 20 s, and then a final extension at 72℃ for 2 min. The staggered primers provide additional sequence complexity for read 2 during sequencing. We purified 400 μL of the first-step PCR with the Zymo DNA Clean & Concentrator and eluted it into 30 μL of elution buffer. We further purified it using a SizeSelect II gel (Invitrogen) to obtain a ∼500 bp product followed by purification and elution using a Zymo DNA Clean & Concentrator column in 20 μL of elution buffer. The purified amplicon was indexed by Q5 polymerase (New England Biolabs) in a 500 μL reaction and a 1 μM equimolar mix of indexing primers for Illumina sequencing. The PCR thermocycling conditions were as follows: initial denaturation at 98℃ for 2 min, followed by 5 cycles of 98℃ for 10 s, 70℃ for 20 s, and then a final extension at 72℃ for 2 min. We purified 100 μL of this second PCR using the Zymo DNA Clean & Concentrator and eluted it into 30 μL of elution buffer. We further purified it using a SizeSelect II gel (Invitrogen), followed by purification and elution in a Zymo DNA Clean & Concentrator column in 20 μL of elution buffer. We quantified them using Qubit dsDNA HS (Invitrogen). We submitted the library to Admera for paired-end sequencing on a lane of the HiSeq 4000 using a custom read1 primer (#75). We used a custom script to associate barcodes with oligos from the read data.

### Pooled editing and plasmid curing

We carried out pooled editing for CRISPEY-BAR+, like previous work (*26*, *29*, *30*). We thawed the yeast library from frozen glycerol stocks and added it to 200 mL SD-histidine-uracil 2% raffinose, starting at an OD600 of 0.4, and incubated at 30℃ with shaking at 250 RPM for 24 h. After this phase of growth in raffinose, we induced editing with galactose. To initiate this new phase, we re-inoculated the pre-editing cultures grown in raffinose in 200 mL SD-uracil 2% galactose, with a starting OD600 of 0.4, and incubated at 30℃ with shaking at 250 RPM for 24 h. This phase of editing induced by galactose lasted for 72 h. We transferred the yeast library to 200 mL of fresh SD-histidine-uracil 2% galactose every 24 hours. We harvested the cells from the last growth in galactose and stored them in SD-histidine-uracil 2% glucose with 15% glycerol at −80℃.

We then removed CRISPEY-BAR+ plasmids after editing by growing them in 200 mL of SD 2% glucose media, starting at an OD600 of 0.4, and incubating at 30℃ with shaking at 250 RPM for 24 h to remove plasmid selection. After this phase of non-selective growth, we inoculated 200 mL SD 2% glucose with 1g/L of 5-fluoroorotic acid monohydrate (5-FOA) (GoldBio), starting at an OD600 of 0.4, and incubated at 30℃ with shaking at 250 RPM for 24 h to select against the *URA*-bearing CRISPEY-BAR+ plasmid (*90*). We froze the edited, plasmid-free yeast library and stored it in 15% glycerol stocks at −80℃ in preparation for the growth competitions.

### Pooled growth competitions

We carried out pooled competitions in 500 mL DeLong flasks (with flat bottoms) in 100 mL of M3+uracil+lysine+leucine media. We thawed edited and plasmid-cured yeast cells (see *Pooled editing and plasmid curing*) in 15 mL of SC media at 30℃, shaking at 250 RPM overnight in a 500 mL flask. We inoculated 100 mL M-uracil-lysine-leucine media starting at an OD600 = 0.4 and grew at 30℃ shaking at 250 RPM for 24h. We then inoculated 500 μL of this saturated culture into 100 mL of fresh media in 500 mL DeLong flasks. This culture was then grown at 30℃ in an incubator sharing at 250 RPM, with incubation time varying depending on the experimental condition: 24 h (1 day), 48 h (2 days), 72 h (3 days), or 120 h (5 days). We included three replicates per condition. After incubation, we transferred 500 μL of the saturated culture into fresh media in a new flask. This serial dilution was continued six times, yielding six time-points over which to measure the rate at which the barcode frequency changed. We pre-warmed fresh media before each transfer. Contamination caused the loss of the last three time-points for one of the three replicates of the 2-day condition. After each transfer of 500 μL, we froze the remaining 9500 μL to sequence the barcodes present at each time-point later. To prepare this culture for freezing, we transferred it to 50 mL conical tubes, spun it down at 3000 RPM for 5 min, resuspended it in 15% glycerol, aliquoted it into three 1.5 mL tubes, and stored it at −80℃.

### DNA extractions

After completing the growth competitions, we extracted DNA from frozen samples using a modified protocol that enhances the extraction yield (*85*). For each sample, a single tube of the three that were frozen for each sample (see *Pooled growth competitions*) was removed from the freezer and thawed at room temperature. We extracted DNA from 400 μL of the sample, and stored the remaining culture at −80℃, using a modified version of the Lucigen MasterPure yeast DNA purification kit (#MPY80200). We transferred the thawed cells into a 2 mL tube and centrifuged at 3500 RPM for 3 min. After discarding the supernatant, the pellet was then resuspended in 400 μL of sterile water and spun again at 3500 RPM for 3 min. After discarding the supernatant, the pellet was resuspended with 1200 μL of the MasterPure lysis buffer supplemented with eight μL of zymolyase and four μL of RNase A (Thermo Scientific). We split the mix into two 1.5 mL centrifuge tubes, which we incubated at 37℃ for 1 hour. After such incubation, we added 2 μL of zymolyase to each tube, and then we incubated them at 65℃ for 4 hours, vortexing the tubes every hour. We then placed the tubes on ice for 10 min and subsequently mixed 300 μL of MPC Protein Reagent with the solution from each tube by vortexing. We separated protein and cell debris by centrifugation at 10,000 RPM for 15 min, transferring 800 μL of supernatant from each tube to a 2 mL centrifuge tube. We further separated the remaining protein and cell debris by centrifuging at 13200 RPM for 5 min. Next, we added 800 μL of isopropanol to each tube, mixed by inversion, and then centrifuged at 10,000 RPM for 15 minutes. The supernatant was discarded. We washed the pellet by adding 500 μL of ethanol 70%, mixing by inversion, and then centrifuging the tubes at 10,000 RPM for 7 min. We discarded the supernatant, and the tubes were left to air-dry. We resuspended the pellet, containing the DNA, in 50 μL of Elution Buffer. Next, we combined the two tubes per sample into a single 1.5 mL DNA LoBind^®^ tube. We added 2 μL of 10 mg/mL RNAse A (Thermo Scientific) and incubated at 37℃ for 30 min to digest the RNA further, and then quantified the amount of DNA using the Qubit dsDNA BR assay (Invitrogen).

### Sequencing library preparation

We amplified 4.5 μg of genomic DNA per sample in a 1200 μL Q5 polymerase (New England Biolabs) reaction with 1 μM forward primer #261 and 1 μM reverse primer equimolar mix of primers #327-#334. We split the PCR reaction into 24 50-μL volumes and performed it with 1 M betaine and initial denaturation at 98℃ for 2 min, followed by 18 cycles of 98℃ for 10 s, 65℃ for 20 s, and then extension at 70℃ for 2 min. The staggered primers provide additional sequence complexity for read2 during the sequencing of the barcode. We purified 200 μL of this first-step PCR with 200 μL of magnetic beads (NucleoMag NGS Clean-up and Size Select, REF 744970.5). The purified amplicons were indexed by Q5 polymerase (New England Biolabs) in a 50 μL reaction and a 1 μM equimolar mix of indexing primers for Illumina sequencing. The PCR thermocycling conditions were as follows: initial denaturation at 98℃ for 2 min, followed by 8 cycles of 98℃ for 10 s, 70℃ for 20 s, then extension at 70℃ for 5 min. We purified the indexed amplicons using 50 μL of beads and eluted them in 100 μL of water. We quantified them using the Qubit dsDNA HS assay (Invitrogen). We mixed equimolar amounts of the purified, indexed amplicons from six time-point samples for each of the three replicates per condition and purified them using a SizeSelect II gel (Invitrogen) to obtain a product of ∼300 bp. We then purified the size-selected libraries with beads. We submitted the libraries to the CZ Biohub SF for paired-end sequencing in a flow cell (4 lanes) of the NovaSeq S4 using a custom read1 primer (#354).

### Read processing

We processed reads in FASTQ format from the competition samples sequenced using NovaSeq using a custom pipeline. We used cutadapt (*91*) to trim adaptors from the reads. We used the following parameters for read2 trimming: 5’ adaptor sequence as GGCCAGTTTAAACTT, 3’ adaptor sequence as `GCATGGC`, maximum error rate of 0.2, and the length of the trimmed read2 as 27 base pairs. We then combined trimmed paired reads using FLASH (*92*) with a minimum overlap of 12 base pairs and a maximum mismatch rate of 0.25. The resulting barcode is 27 base pairs in length. The barcodes with a perfect match to those barcodes from the barcode-oligo map were counted for analysis as described below.

### Estimating edited variant effect on fitness

Incorporating multiple barcodes per edit enhances the precision of fitness measurements by increasing biological replication within a flask. This approach enables the detection of random mutants that may arise during transformation, editing, or competition, as well as cases where the specific edit failed in the CRISPEY-BAR+ procedure, which appear as outliers, thereby improving the accuracy of our estimates of variant fitness effects. Spontaneous mutations with strong positive fitness effects, in particular, can disproportionately dominate the reads for a given variant. However, these mutations are unlikely to impact multiple independently edited barcodes. To mitigate false positives, we removed these outlier barcodes by discarding the barcode with the highest read count for each variant. We aggregated all the barcodes for the same edit to infer fitness from the edit count trajectories.

To infer fitness from the edit count trajectories, we followed a variational Bayesian approach (*52*). Briefly, our objective is to infer the fitness values *s*^≥^ = (*s*^(1)^, *s*^(2)^, …, *s*^(N)^) for all *N* edited strains. For a specific strain *j*, our data consists of a vector of barcode counts across all time points sequenced in the experiment,

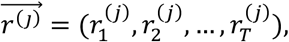

for the *T* time points in our experiment. The entire dataset then consists of *N* vectors of raw reads

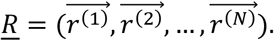

Our fitness models for how the allele frequency of barcode *j* changes over time takes the form

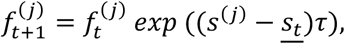

where *f_t_^(j)^* is the allele frequency of barcode *j* at time *t*, <u>*s*_t_</u> is the population mean fitness at time *t*, and τ is the time interval between *t* and *t* + 1. Thus, we must include the inference of the nuisance parameter <u>*s*_)_</u> for each time point in our experiment. To constrain the value of this parameter, the experiment includes a series of what we consider to be neutral barcodes (barcodes associated with non-editing oligos) whose fitness is zero by definition. This means that we include a series of barcodes that, by definition, have

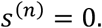

Furthermore, the barcode frequency *f_t_(j)* is not what our experiments quantify. Naively, we could simply assume that

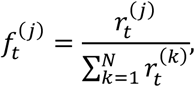

but this would assume we have perfect readouts for the experiment. Since we use a Bayesian framework, we can simply include the computation of all frequencies

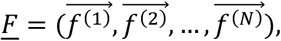

as nuisance parameters in our inference.

By Bayes theorem, our inference problem can be expressed as

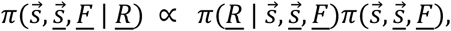

where <u>*s*</u> = (*s*<u>_1_</u>, *s*<u>_2_</u>, …, *s*<u>_N_</u>), and π(⋅) is a probability distribution. Although this is an intractable problem, Razo-Mejia *et al.* (*52*) deploy a variational approach to solve this inverse problem of extracting fitness values from raw read counts.

Furthermore, using a hierarchical Bayesian model, the Bayesian framework can be extended to combine the information from multiple experimental replicates into a single inference problem. Hierarchical models systematically account for all available information and propagate sources of uncertainty throughout the inference process, increasing the statistical power when inferring the fitness values. The results shown in the main text take this hierarchical approach as follows: The reads for all individual barcodes mapping to a specific edit were aggregated to be used as a single number *r*^(j,n)^_t_ for edit *j* and replicate *n* at time *t*. We then assume that we can infer the corresponding fitness *s*^(’,+)^ as a sample from a hyper-parameter distribution (the hyperfitness) Φ^(j)^ shared among all edits *j* across all replicates. The objective then becomes inferring the values for all hyperfitness values Φ = (Φ^(1)^, Φ^(2)^, …, Φ^(N)^) for all *E* edits and the individual replicate fitness values *s* = (*s*^(1)^, *s*^(2)^, …, *s*^(N)^) for all *M* experimental replicates given the corresponding raw reads *R* = r^(1)^, …, *r^(M)^*. This inference problem can be written by Bayes theorem as (we ignore the corresponding nuisance parameters for simplicity although they are included in the full inference; see (*49*) for more details)

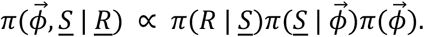

In this way, all experiments are connected through the hyperfitness parameter, which brings together all information from each replicate and its corresponding uncertainty. The main text results are based on the posterior distributions of the hyperfitness parameters.

### Variant properties

We annotated variants using Variant Effect Predictor (VEP) (*93*). We retrieved the predicted protein structures of all *S. cerevisiae* S288C genes in Supplementary Table 1 from the AlphaFold Protein Structure Database (*94*, *95*). There are no protein structures for *IRA1*, *IRA2*, and *SDC25*. We measured solvent accessibility using DSSP (*96*, *97*). We measured packing density as the number of other α-carbons within 10Å of the α-carbon for a given residue using custom code. We used FoldX to determine the destabilizing effect of nonsynonymous variants (*58*). For all yeast genes, we retrieved SIFT scores from the mutfunc database (*98*). We obtained the allele frequency and heterozygosity of each variant from the 1,011 yeast genomes collection (*20*). From the same source, we received the number of singleton variants for each of the 1,011 strains and their ploidy. For a given singleton variant, we use the number of other singleton variants present in the same strain as a proxy for variant age, following Zhu *et al.* (*59*).

While this metric serves as a reasonable proxy on average—since under a strict molecular clock with constant effective population size (*Nₑ*), branch length reflects divergence time—variation in *Nₑ* can introduce noise. Lineages with smaller Nₑ may accumulate more slightly deleterious mutations, leading to longer apparent branch lengths than lineages with larger *Nₑ*, even under identical divergence times.

### Constructing a maximum likelihood phylogenetic tree

To construct a maximum likelihood phylogenetic tree for the 1,011 natural isolates we used the published whole-genome sequencing data from Peters *et al.* (*20*). We started with the previously published VCF file, which contains every single-nucleotide polymorphism (SNP) and indel that has been called at the population level (1011Matrix.gcvf). We filtered for SNP sites present in all strains and where the minor allele could be confidently called in at least 2 strains and could thus be parsimony informative. A maximum likelihood tree was generated based on the 866,130 filtered SNPs using IQTREE2 (iqtree2-s filtered_snps.min4.phy -m GTR+ASC) (*99*). The resulting maximum likelihood tree was rooted using the Taiwanese clade as an outgroup as this represented the most divergent population (*20*). This rooted tree was used in all subsequent analyses (Fig. 1B and Fig. 2A).

### Variant co-occurrence analysis

For each pair of alleles (*i* and *j*) the probability of co-occurring in the same natural isolate was calculated as the Jaccard similarity coefficient:

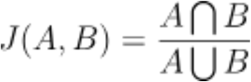

where *A* is the set of natural isolates in which allele *i* is present and *B* is the set of natural isolates in which allele *j* is present. Using this measure, we quantified whether pairs of non-neutral variants were significantly less likely to co-occur than pairs of neutral variants as this would suggest that they were under purifying selection (since a deleterious variant is expected to go extinct before a second deleterious variant can arise in the same genetic background).

We calculated the mean Jaccard similarity for all pairs of singleton non-neutral variants and then compared this with the mean Jaccard similarity for all pairs of neutral singleton variants (Fig. 4F). To determine whether the difference in means was statistically significant, we repeated this calculation for 1,000 bootstrap replicates. The *P*-values in Figure 4F corresponds to the proportion of bootstrap replicates in which neutral variants co-occurred less frequently than the non-neutral variants considered.

### Identifying exclusively natural, exclusively industrial, and clade-exclusive variants

Of the 1,011 natural isolates reported in Peters *et al.* (*20*), 963 isolates belong to one of 29 distinct clades (Fig. 2A), and were assigned to one of 23 different ecological origins in that study. We grouped these ecological origins into three categories: industrial, natural, and other (Fig. 1B). Specifically, 587 strains were classified as “industrial” if their ecological origin was listed as either beer, bakery, industrial, wine, distillery, palm wine, bioethanol, fermentation, cider, dairy or sake. We classified 392 strains as “natural” if their ecological origin was listed as either water, fruit, flower, human-clinical, insect, nature, tree, soil, or human. Finally, 32 strains were classified as “other” if their ecological origin was listed as either laboratory strains, probiotics, or unknown. Strains in this last category were excluded from subsequent analyses.

Using the VCF file from Peters *et al.* (*20*), we determined the ecological and clade-level distribution of 9,447 variants from the Ras/PKA and TOR/Sch9 pathways for which we could estimate selection coefficients in our pooled competition assays. Of these variants, 2,851 were observed exclusively in industrial strains, whereas 3,250 were exclusive to natural strains. At the clade level, 5,778 variants were restricted to a single clade, whereas 2,994 were present in multiple clades. The remaining 675 variants were found in the 48 strains without clade assignments and were treated as a single group in clade-level analyses (see *Quantifying variant enrichment in industrial strains*).

### Quantifying the enrichment of beneficial variants in industrial strains

We determined whether beneficial variants were disproportionately enriched in industrial versus natural strains using three complementary statistics each capturing variant enrichment at different phylogenetic scales. First, as a straightforward measure of enrichment, we calculated the odds ratio (OR) comparing the likelihood that an exclusively industrial variant was beneficial to that of a solely natural variant (Fig. 5A). Second, we computed the Spearman’s rank correlation (ρ) between the proportion of clade-exclusive variants that were beneficial and the fraction of industrial strains in each clade, to assess how enrichment in beneficial variants tracks phylogenetic structure (Fig. 5B). Third, we fit a generalized linear mixed-effects model (GLMM) with clade as a random effect to estimate the association between ecological origin and the probability of being beneficial while controlling for shared ancestry.

For the OR, statistical significance and 95% confidence intervals were obtained using the *fisher.test* function in R. For ρ, we calculated the correlation between (a) the proportion of clade-exclusive variants that were beneficial and (b) the proportion of industrial strains in each clade. To account for sampling uncertainty, we generated 10,000 bootstrap replicates by resampling clades with replacement, weighting each by the number of variants it contained. The bootstrap distribution was used to derive 95% confidence intervals and p-values. For the GLMM, we fit a binomial model of the form *IsBeneficial ∼ Type_Eco + (1 | Clade)* using the *lme4* package in *R*. The fixed-effect estimate for the “industrial” category gives the log-odds difference in the probability of a variant being beneficial relative to exclusively natural variants, accounting for clade-level differences. 95% confidence intervals were computed using the *confint* function, and *P*-values were obtained from the model summary.

We repeated the above analysis for deleterious non-neutral variants and never observed any statistically significant enrichment in industrial strains for any of our three statistics (results for the odds ratio are also shown in Fig. 5A). We also repeated the above analysis separately for variants found within one of the two pathways considered (Fig. S10A), and for variants in each of the 24 individual genes we have studied (Fig. S10B).

### Permutation tests controlling for allele frequency and functional impact

To test whether the enrichment of beneficial Ras/PKA pathway variants in industrial strains could be explained by differences in allele frequency or predicted functional impact (SIFT score), we performed permutation analyses matched on these properties. Specifically, we binned all variants into 20 roughly equal-width bins based on log10-transformed allele frequencies, or separately into 20 bins based on their SIFT scores (for nonsynonymous variants only). Within each bin, we permuted the beneficial status of variants without replacement, thereby preserving the original distribution of allele frequencies or SIFT scores while randomizing whether or not a given variant is beneficial or not.

For each of 10,000 permutations, we recalculated the three statistics described above— odds ratio, clade-level Spearman correlation, and GLMM effect size—providing a null distribution under the hypothesis that enrichment arises solely from differences in allele frequency or predicted functional impact. The observed statistics were then compared to the permutation distributions to assess whether they could be explained by these variant properties alone (Fig. S9A-F).

To test whether the observed enrichment of beneficial variants in industrial strains was specific to Ras/PKA and TOR/Sch9 pathways or could be explained by genome-wide effects such as relaxed purifying selection due to domestication bottlenecks, we performed permutation tests using matched control sets of non-synonymous variants outside these pathways. We first identified non-synonymous variants exclusive to industrial or natural strains, excluding those missing SIFT scores. Variants within the Ras/PKA and TOR/Sch9 pathways (“CRISPEY” variants) were flagged, and their beneficial status was assigned based on the CRISPEY screen results.

To generate genome-wide controls, we matched each non-synonymous CRISPEY variant to one or more genome-wide non-synonymous variants with identical SIFT score and log10-transformed allele frequency bins (20 bins each). For each of 10,000 permutations, we randomly sampled one matched variant per CRISPEY variant, assigning it the original beneficial status of the corresponding CRISPEY variant. We then recalculated the odds ratio, clade-level Spearman correlation, and GLMM effect size for matched controls to obtain null distributions that account for differences in allele frequency and predicted functional impact across the genome (Fig. S9G-I).

---

**Figure S1.**
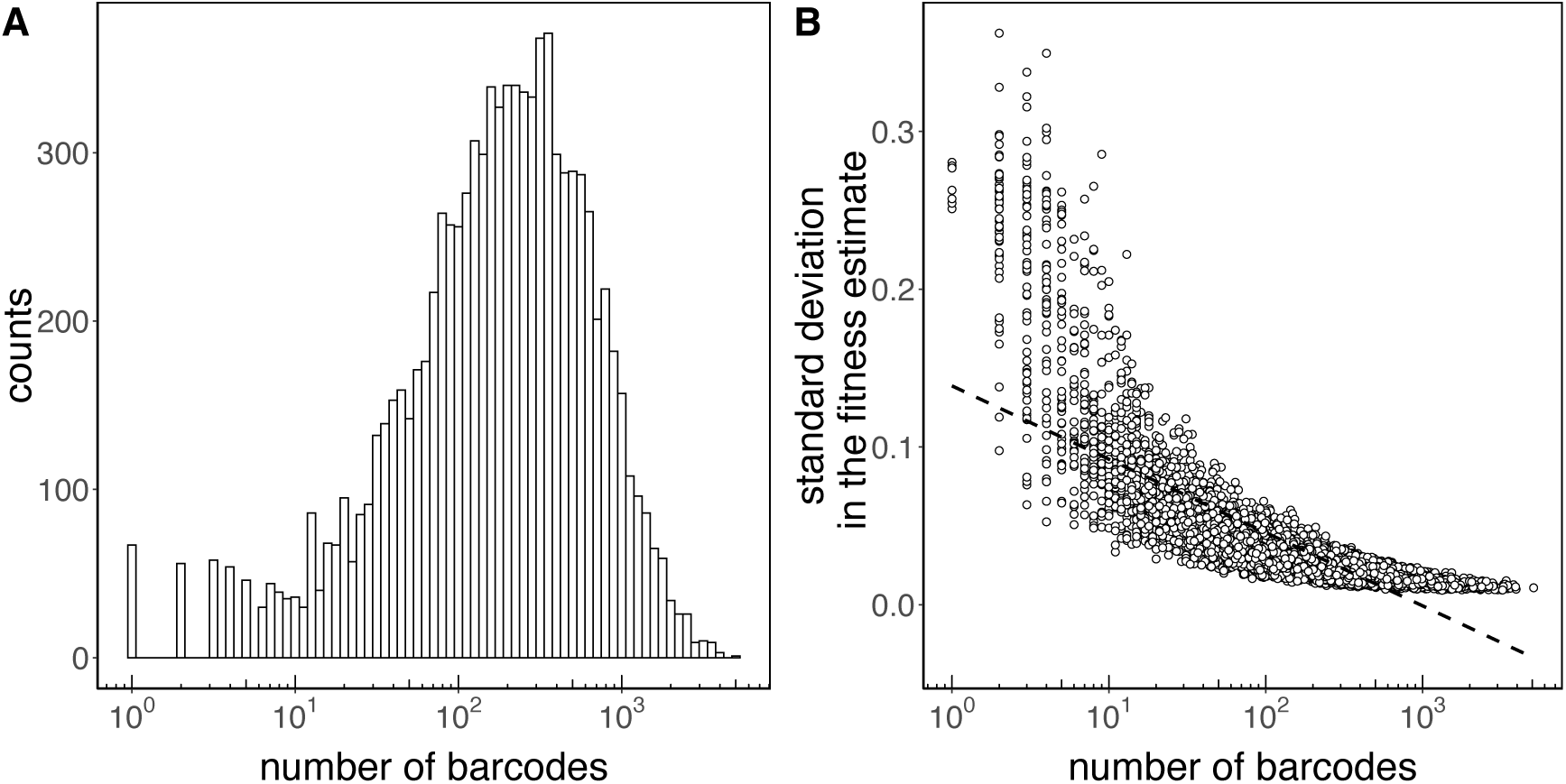
Barcode coverage and its relationship to measurement precision. (A) Histogram showing the number of barcodes per edit for all edits at timepoint = 0 of the pooled growth competitions (Methods). (B) For the same edits, standard deviation of the fitness estimate in the 2D condition as a function of the number of barcodes. A dashed line shows the best least-squares fit to the data provided as a visual guide. The x-axis is displayed on a log_10_ scale in both panels. See Methods and the first section of Results for details on pooled growth competitions, experimental conditions, and fitness estimation.

**Figure S2.**
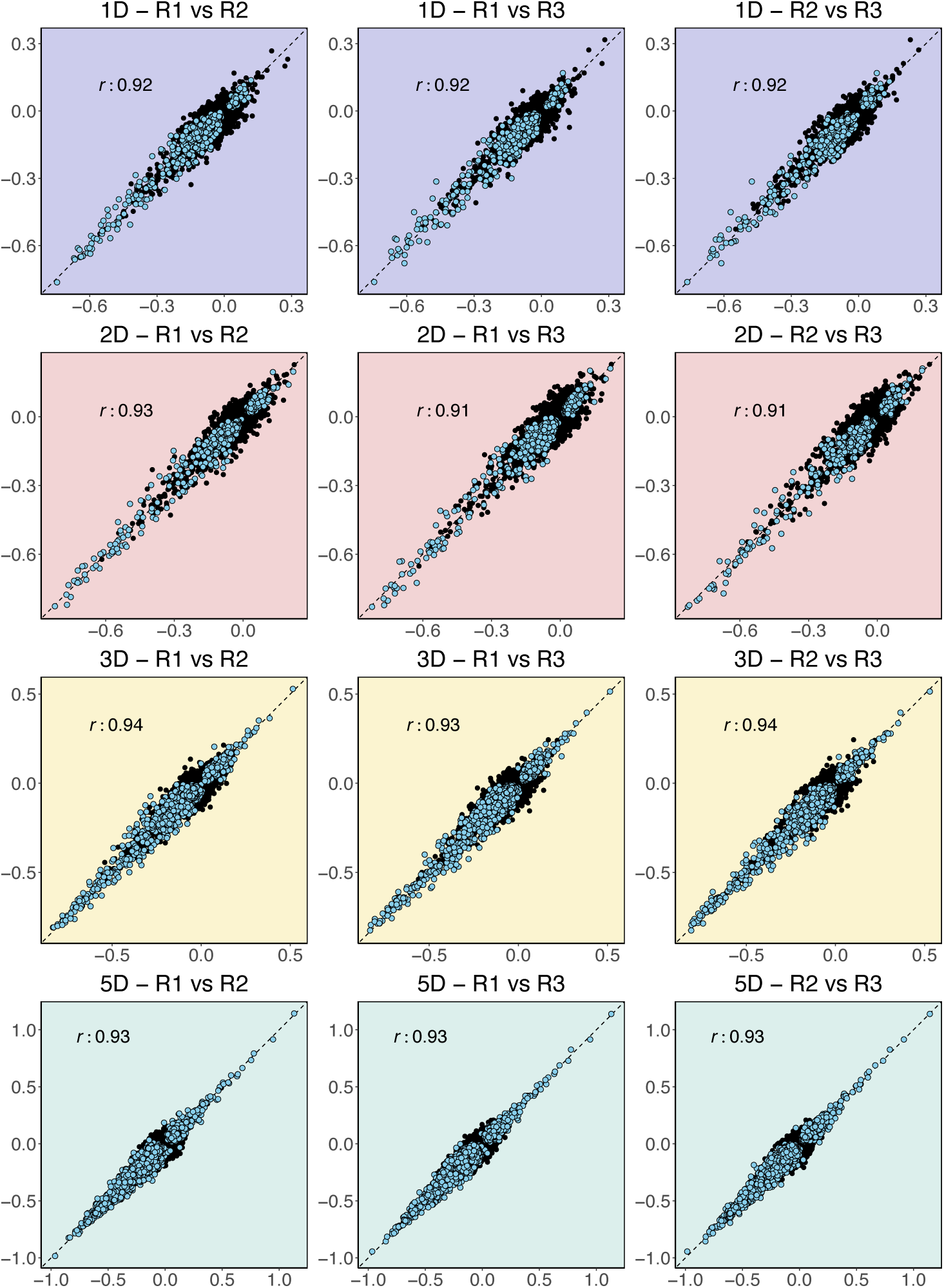
Fitness estimates are highly reproducible across technical replicates in the four conditions. Scatter plots show pairwise comparisons of fitness values for each variant across replicates (R1, R2, and R3) in each condition (1D, 2D, 3D, 5D). Neutral variants are shown in black, while non-neutral variants—those with significant fitness effects—are highlighted in blue. Dashed diagonal lines indicate the identity line, and Pearson’s correlation coefficients (*r*) quantify the agreement between replicates.

**Figure S3.**
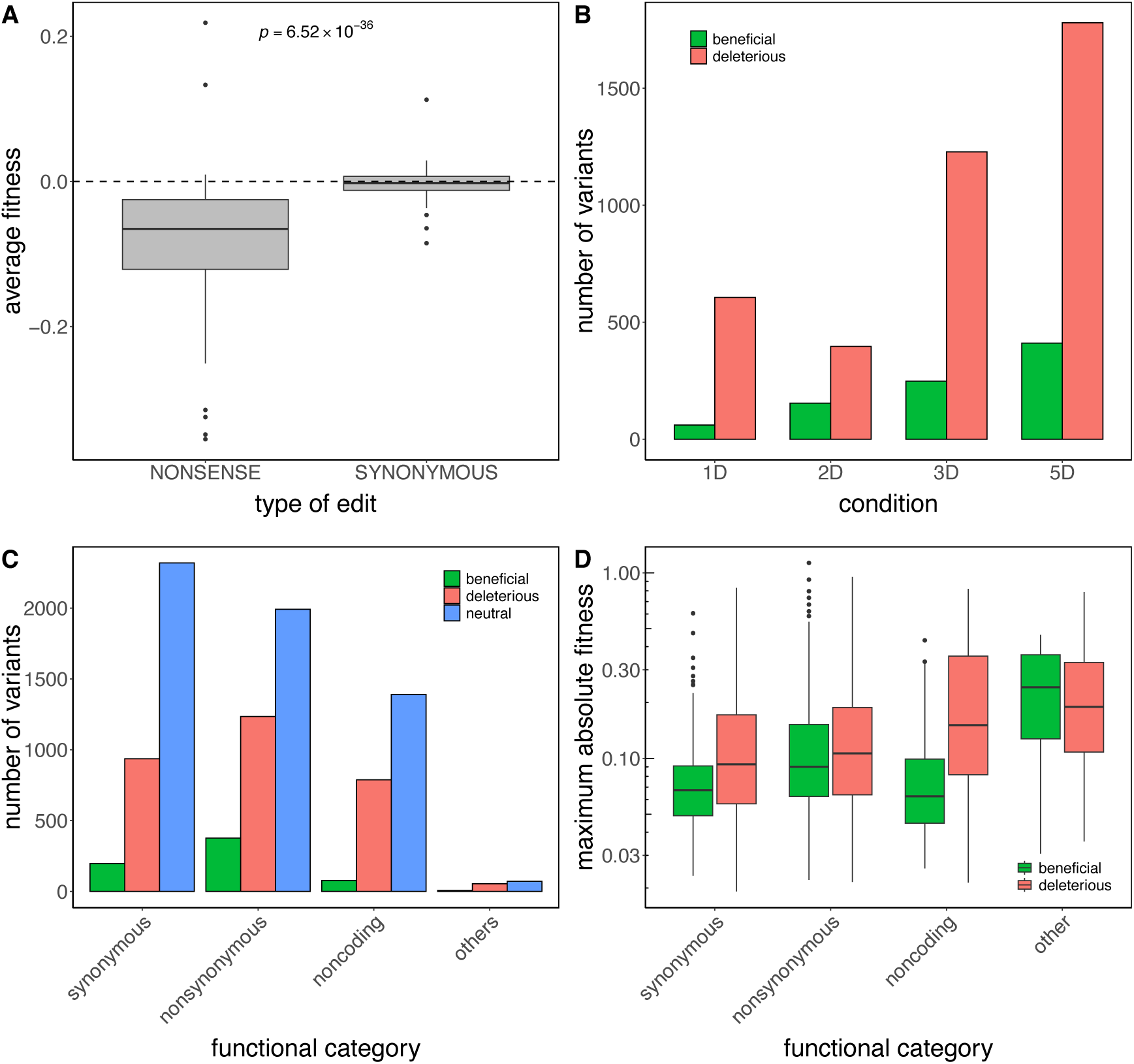
(A) Box plot of average fitness effects across the four tested conditions for edits modifying cysteine or tyrosine codons into stop codons, and edits introducing synonymous substitutions in the same codons. (B) Bar plot of the number of beneficial and deleterious variants by experimental condition (1-Day condition, 2-Day condition, 3-Day condition, and 5-Day condition). (C) Bar plot of the number of neutral, deleterious, and beneficial variants by variant effect. (D) Box plot of the maximum absolute value of fitness effects across the four conditions tested by variant effect for beneficial and deleterious variants.

**Figure S4.**
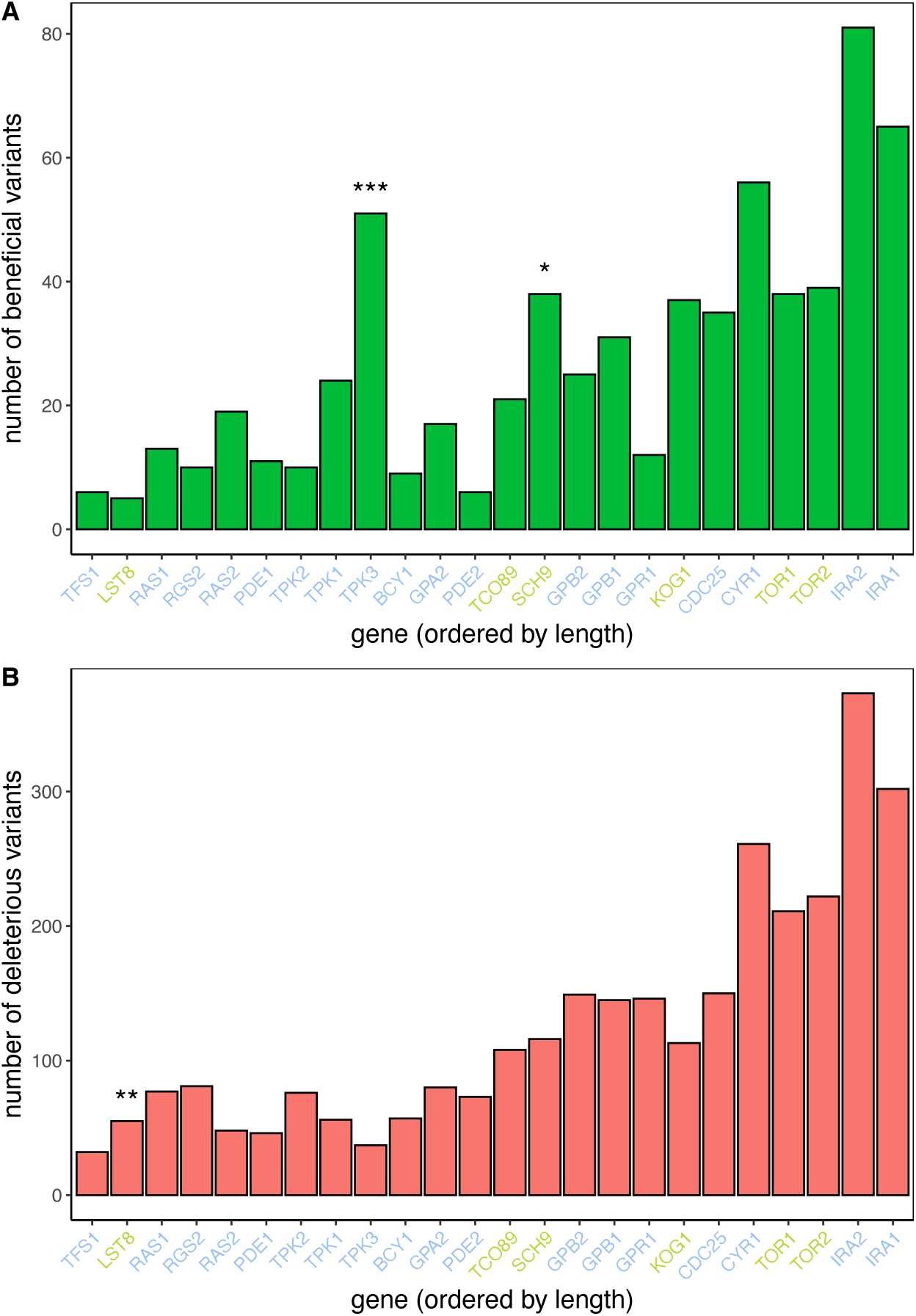
Variant burden across signaling genes in the Ras/PKA and Tor/Sch9 pathways. Bar plots show the number of beneficial (panel A) and deleterious (panel B) variants per gene, ordered by increasing gene length. Bars are colored by variant type: green for beneficial, and red for deleterious variants. Gene names are colored by pathway: blue for Ras/PKA and green for Tor/Sch9 (following the color scheme in Fig. 2E). Asterisks indicate genes with a statistically significant excess of variants based on a binomial test with multiple-testing correction applied at the pathway level (*FDR < 0.05; **FDR < 0.01; ***FDR < 0.001).

**Figure S5.**
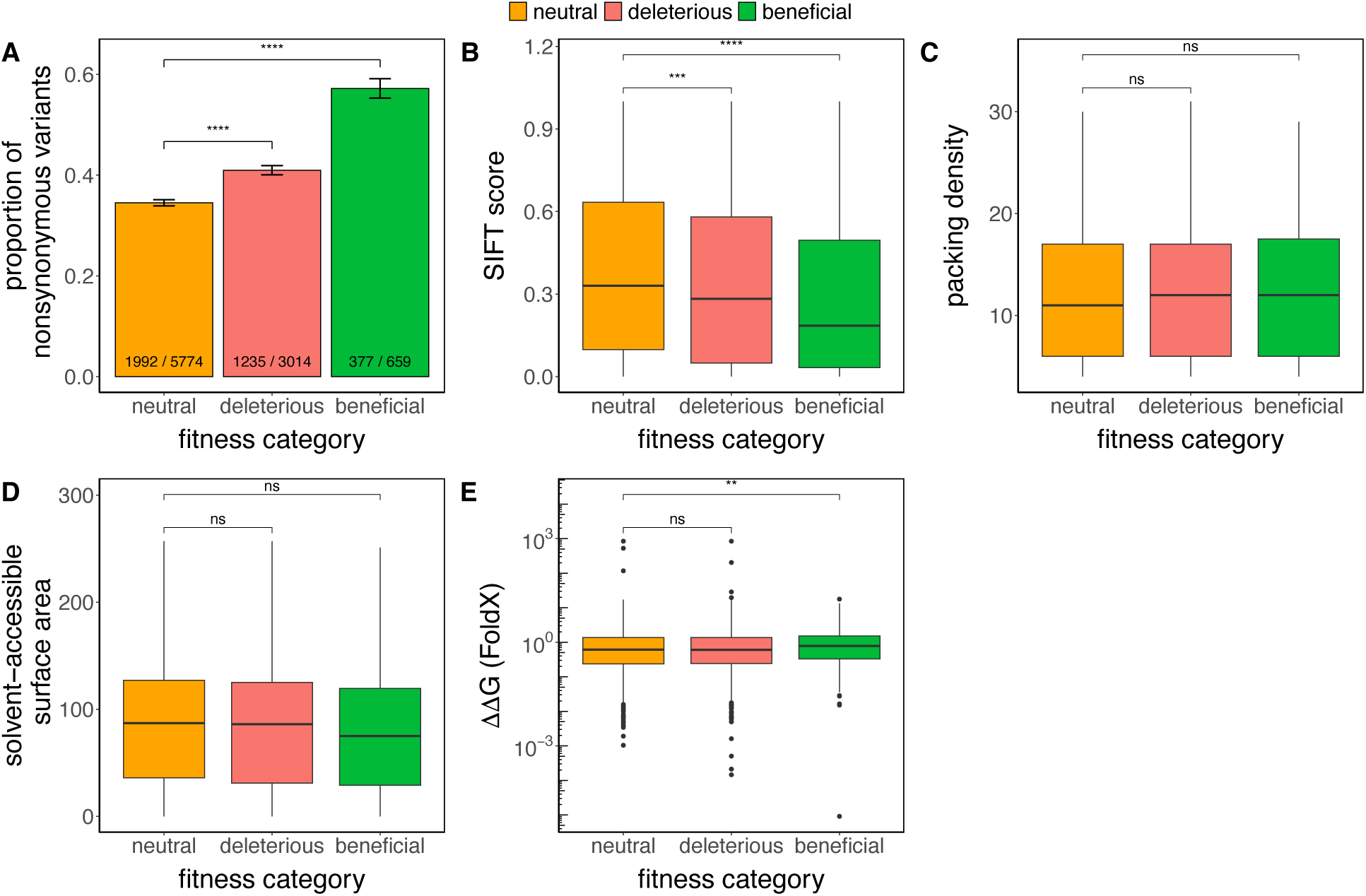
To accompany Figure 3. Non-neutral variants tend to be missense and are predicted to be functional. (A) Bar plot showing an enrichment of nonsynonymous variants among non-neutral variants compared to neutral variants. In this and all subsequent panels, non-neutral variants are grouped as beneficial if they have a positive fitness effect in at least one condition, or as deleterious otherwise. The ratios indicate the number of nonsynonymous variants among all the variants of each class. (B) Box plot of SIFT scores for neutral, deleterious, and beneficial variants. (C) Box plot of packing density for neutral, deleterious, and beneficial variants. (D) Box plot of solvent accessibility for neutral, deleterious, and beneficial variants. (E) Box plot of ΔΔG as computed by FoldX for neutral, deleterious, and beneficial variants. Differences between non-neutral and neutral variants were assessed using Fisher’s exact test for panel A, and the Wilcoxon test for the remaining panels. Significance levels are indicated as follows: *P* ≤ 0.0001: ****, *P* ≤ 0.001: ***, *P* ≤ 0.01: **, *P* ≤ 0.05: *, not significant: ns. Box plots show the median an interquartile range; whiskers extend to 1.5 times the interquartile range.

**Figure S6.**
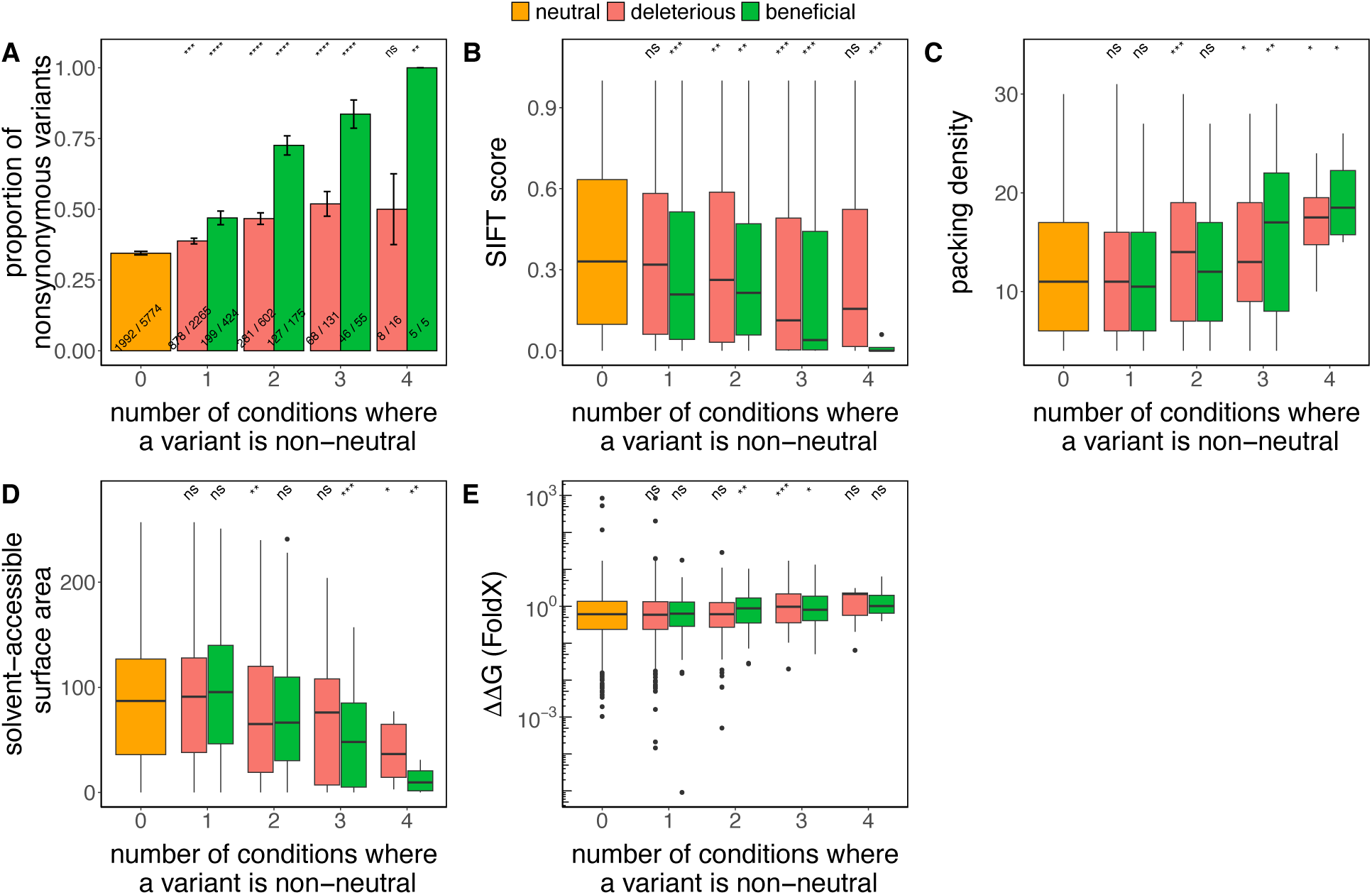
To accompany Figure 3. Non-neutral variants tend to be missense and are predicted to be functional. (A) Bar plot showing an enrichment of nonsynonymous variants among non-neutral variants compared to neutral variants. In this and all subsequent panels, non-neutral variants are grouped as beneficial if they have a positive fitness effect in at least one condition, or as deleterious otherwise, and by the number of conditions in which they exhibit non-neutral fitness effects. The ratios indicate the number of nonsynonymous variants among all the variants of each class. (B) Box plot of SIFT scores for neutral, deleterious, and beneficial variants. (C) Box plot of packing density for neutral, deleterious, and beneficial variants. (D) Box plot of solvent accessibility for neutral, deleterious, and beneficial variants. (E) Box plot of ΔΔG as computed by FoldX for neutral, deleterious, and beneficial variants. Differences between non-neutral and neutral variants were assessed using Fisher’s exact test for panel A, and the Wilcoxon test for the remaining panels. Significance levels are indicated as follows: *P* ≤ 0.0001: ****, *P* ≤ 0.001: ***, *P* ≤ 0.01: **, *P* ≤ 0.05: *, not significant: ns. Box plots show the median an interquartile range; whiskers extend to 1.5 times the interquartile range.

**Figure S7.**
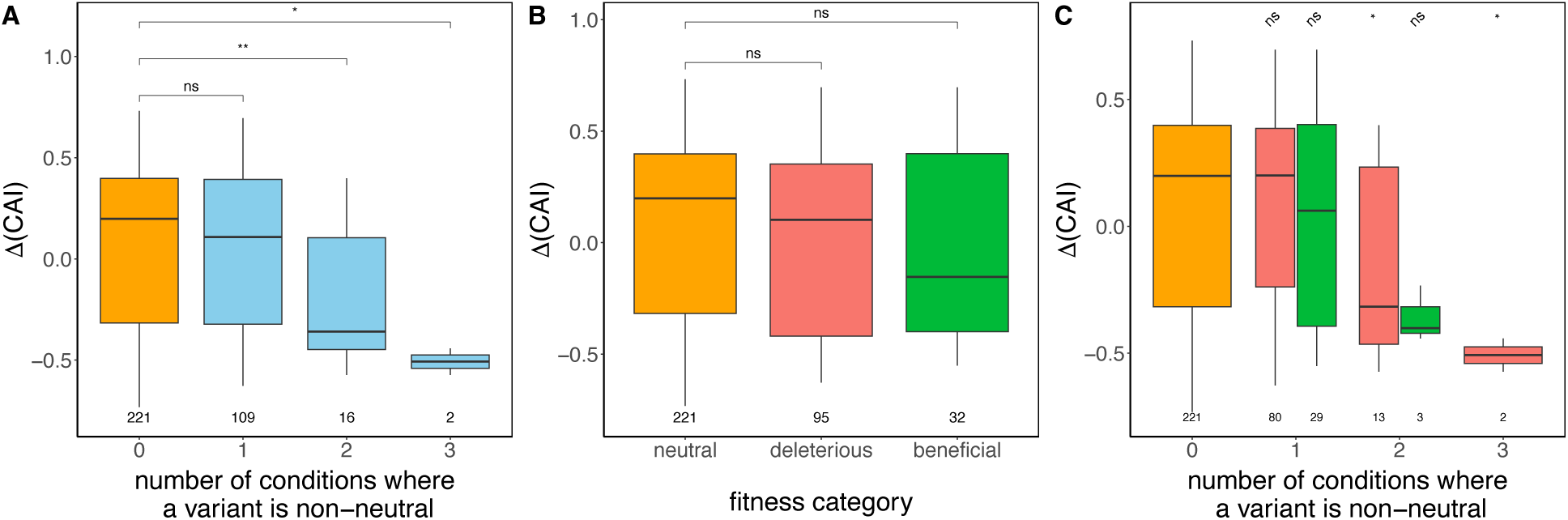
Synonymous variants with fitness effects tend to decrease the codon adaptation index (CAI) when located near the *N*-terminus. (A) Box plots showing the change in codon adaptation index (ΔCAI) between reference and variant codons for synonymous variants located within the first 10% of the protein. Variants are grouped by the number of experimental conditions in which they exhibit significant non-neutral fitness effects. (B) The same ΔCAI values for these variants are now grouped by fitness effect category (beneficial, neutral, or deleterious). (C) ΔCAI values for the same synonymous variants, grouped by both fitness effect category and the number of conditions in which the variant is non-neutral. Box plot center lines show medians, boxes indicate interquartile ranges, and whiskers extend 1.5 times the interquartile range. Sample sizes are indicated below each group. Statistical significance was assessed using pairwise Wilcoxon rank-sum tests; P values are shown as follows: *P* ≤ 0.01: **, *P* ≤ 0.05: *, not significant: ns.

**Figure S8.**
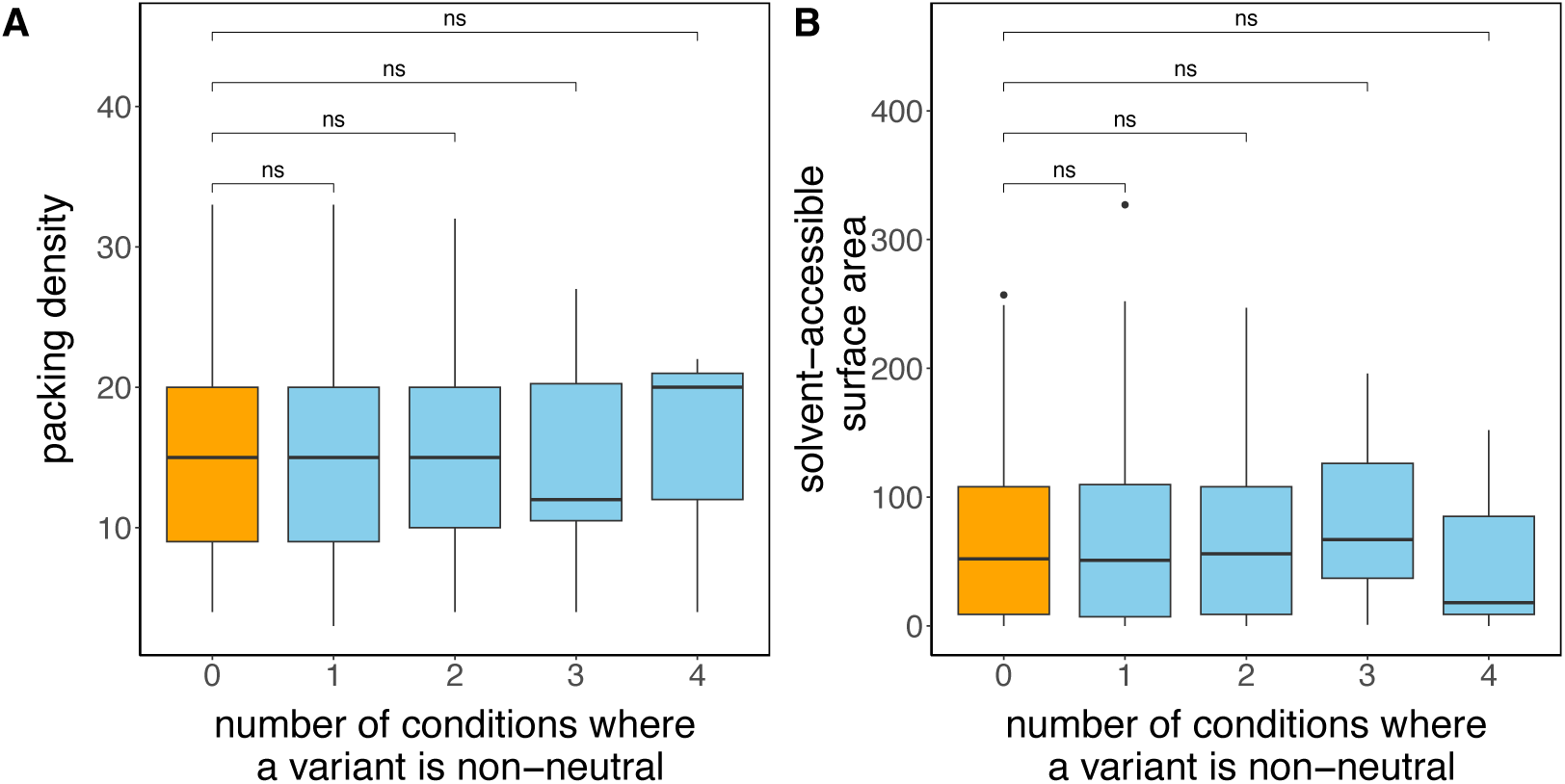
Structural context does not explain synonymous fitness effects. (A) Residue packing density and (B) solvent-accessible surface area (SASA) at codons harboring synonymous variants, stratified by the number of conditions in which each variant is non-neutral (0–4). Orange boxes indicate variants neutral in all conditions (0); blue boxes indicate variants non-neutral in at least one condition. Brackets show pairwise Wilcoxon rank-sum tests: “ns” denotes *P* > 0.05. Box plots show the median and interquartile range; whiskers extend to 1.5 times the interquartile range.

**Figure S9.**
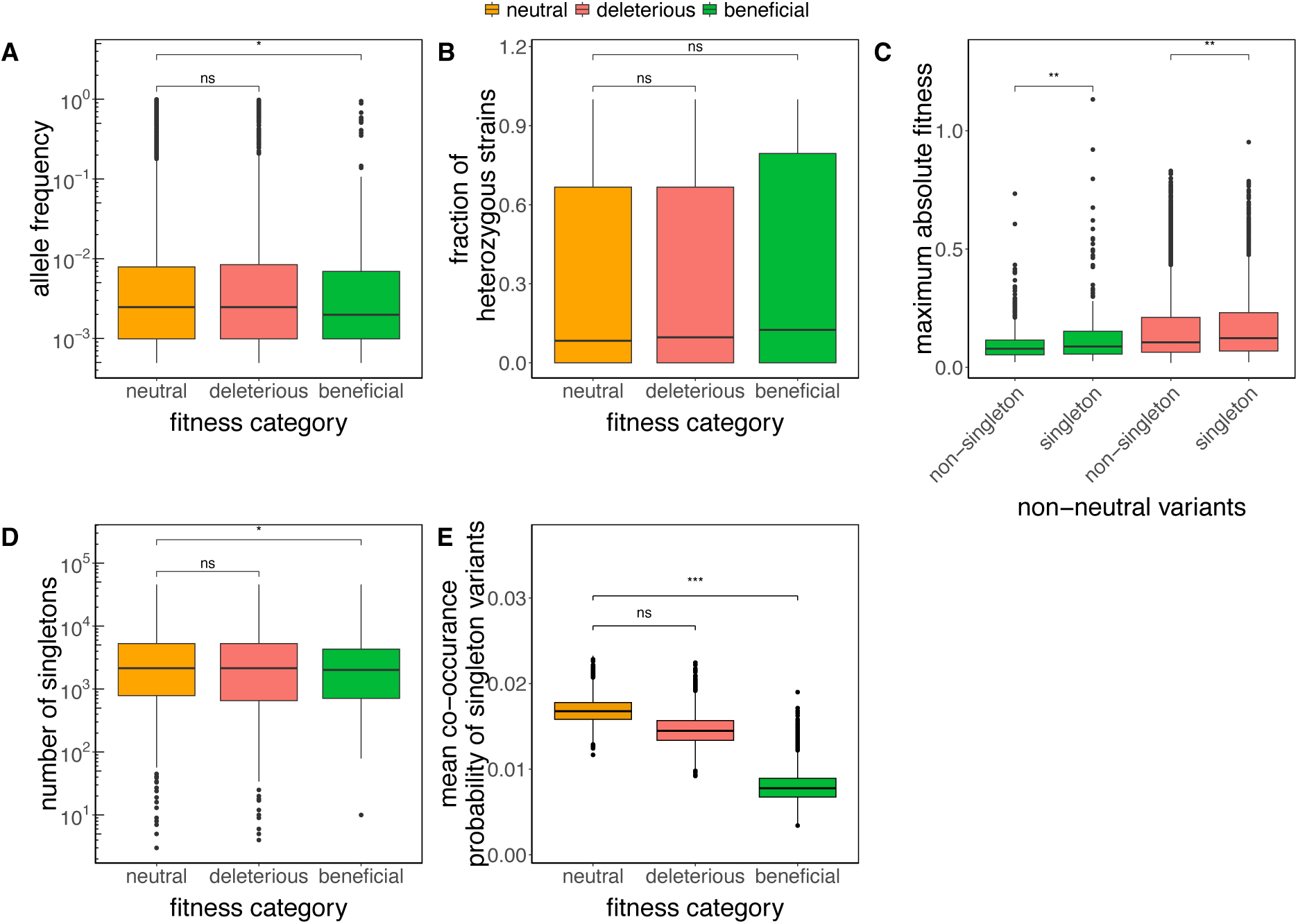
To accompany Figure 4. Low-frequency variants show evidence of purifying selection. (A) Box plot of allele frequency for neutral, beneficial, and deleterious variants. In this and all subsequent panels, non-neutral variants are grouped as beneficial if they have a positive fitness effect in at least one condition, or as deleterious otherwise. (B) Box plot showing the fraction of strains where the variant is in heterozygosis for neutral, beneficial, and deleterious variants. (C) Box plot showing that the experimental fitness impact of rare variants is greater than for more common variants for both beneficial (in green) and deleterious variants (in red). Rare variants are defined as those present in a single strain (singletons). (D) Box plot of the age of a variant for neutral, beneficial, and deleterious variants. (E) Box plot showing the co-occurrence probability of singleton variants by fitness category. The data shown represents the mean co-occurrence for 10,000 bootstrap samples (Methods). Significance levels are indicated as follows: *P* ≤ 0.0001: ****, *P* ≤ 0.001: ***, *P* ≤ 0.01: **, *P* ≤ 0.05: *, not significant: ns. Differences between groups were assessed using a Wilcoxon’s test. Box plots show the median an interquartile range; whiskers extend to 1.5 times the interquartile range.

**Figure S10.**
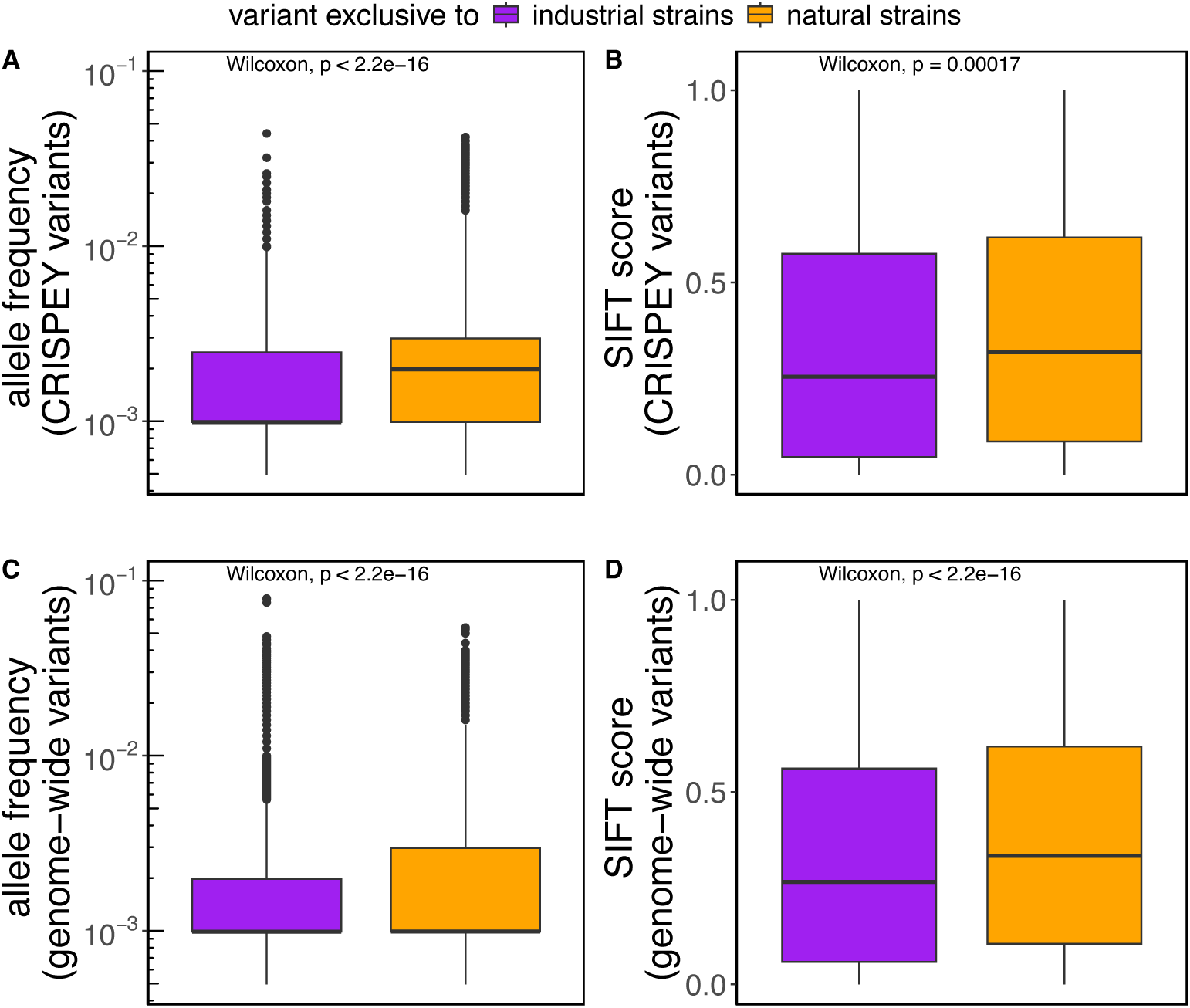
Comparison of allele frequency and predicted functional impact (SIFT score) for variants exclusive to industrial or natural strains. (A-B) Box plots showing differences in allele frequency (A) and SIFT score (B) for CRISPEY variants exclusively found in industrial or natural strains. (C–D) Box plots showing differences in allele frequency (C) and SIFT score (F) for nonsynonymous variants outside of the 24 Ras/PKA and TOR/Sch9 genes we have studied. Box plots show the median and upper and lower quartiles; whiskers show 1.5 times the interquartile range. *P*-values are from Wilcoxon rank-sum tests.

**Figure S11.**
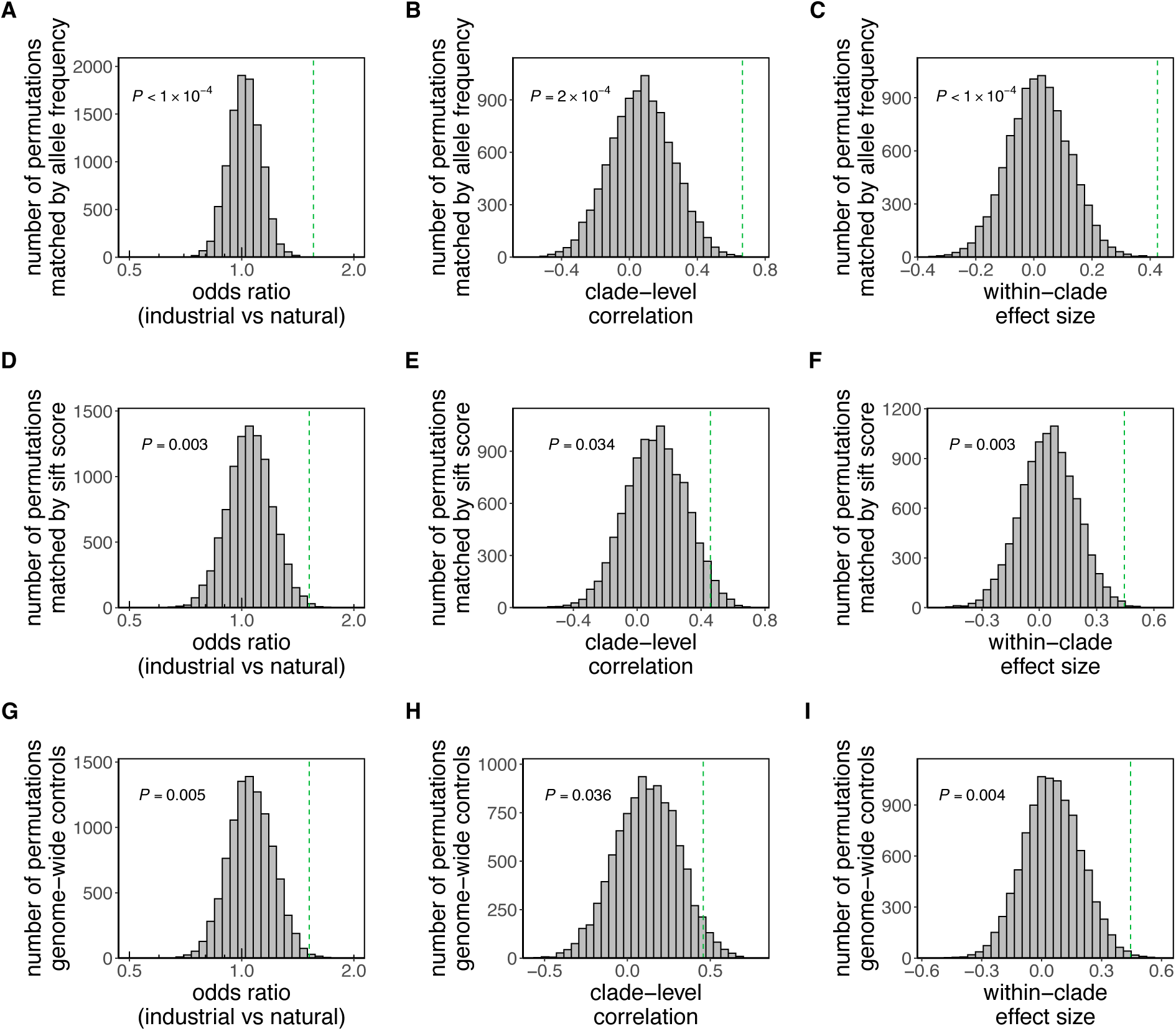
Permutation-based analysis of the significance of enrichment in beneficial variants in industrial strains, accounting for possible confounders. (A–C) Null distributions from allele-frequency–matched permutations of the measured variants, showing observed odds ratio (A), clade-level correlation (B), and within-clade model effect size from a mixed-effect logistic regression (Methods) (C). (D–F) Same as A–C, but with permutations matched on SIFT score for non-synonymous variants. (G–I) Same as A–C, but with genome-wide nonsynonymous variants (outside studied-targeted genes) as controls matched by allele frequency and SIFT score (Methods). Dashed vertical lines indicate observed enrichment; *P*-values reflect the fraction of permutations exceeding the observed enrichment.

**Figure S12.**
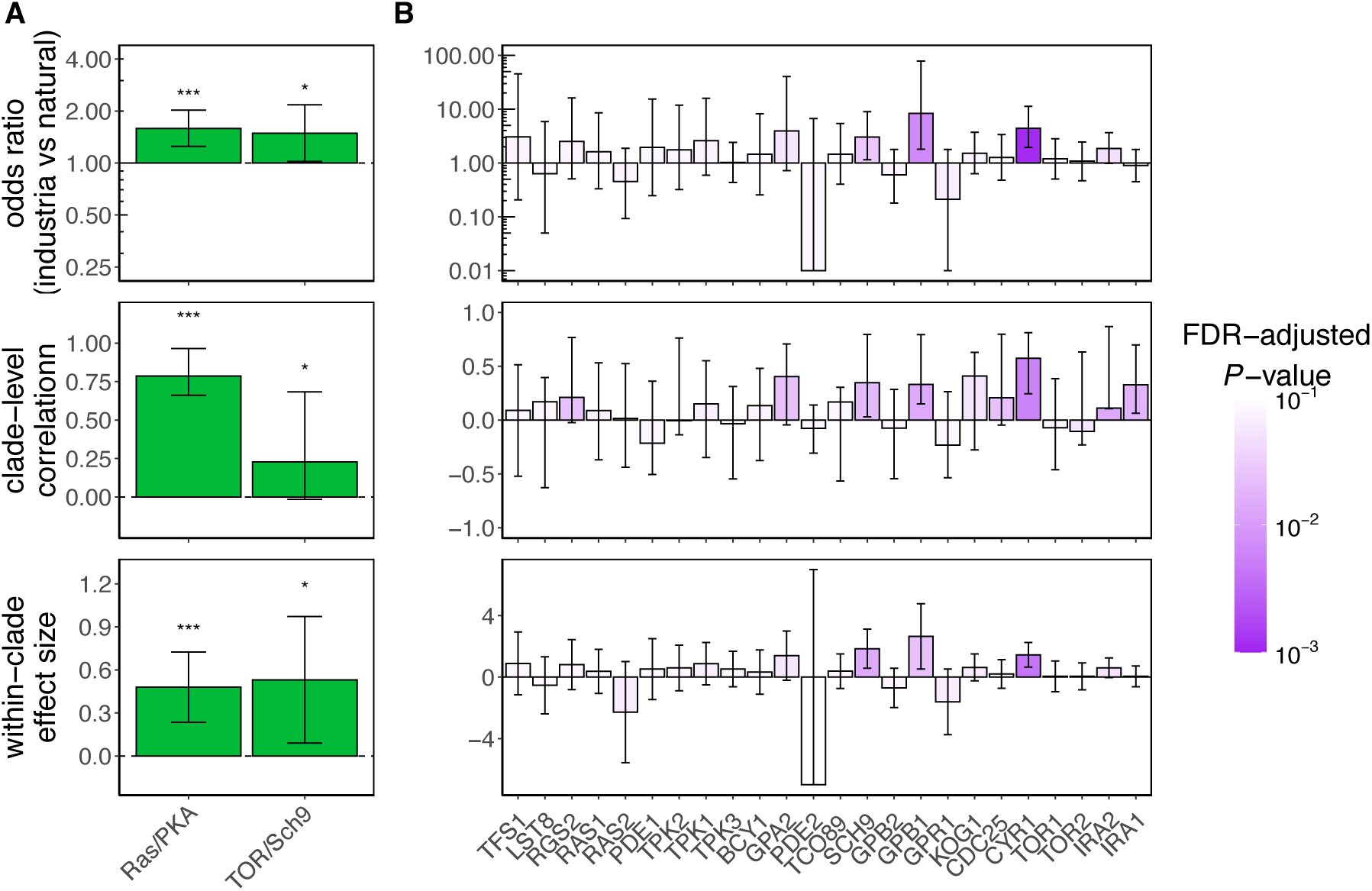
Pathway and gene-level analysis of beneficial variant enrichment in industrial strains. (A) Pathway-level analysis showing (from top to bottom) the odds ratio of beneficial variant enrichment in industrial strains relative to natural strains, the clade-level correlation (Spearman’s ρ), and the within-clade effect size from a mixed-effect logistic regression (Methods). Error bars represent 95% confidence intervals estimated via bootstrapping (Methods). Significance levels are indicated by asterisks (\**P* < 0.05; \*\**P* < 0.01; \*\*\**P* < 0.001; ns = not significant). The horizontal dashed line indicates the null expectation (odds ratio = 1 for enrichment, correlation = 0, effect size = 0). (B) Gene-level analysis showing odds ratios of beneficial variant enrichment (top), clade-level correlations (middle), and within-clade effect sizes (bottom). Bars are colored by log10-transformed FDR-adjusted p-values, with error bars indicating 95% confidence intervals (Methods). Horizontal dashed lines indicate null expectations as in (A). Gene names are ordered by increasing protein length and displayed on the x-axis with labels rotated for readability.

**Figure S13.**
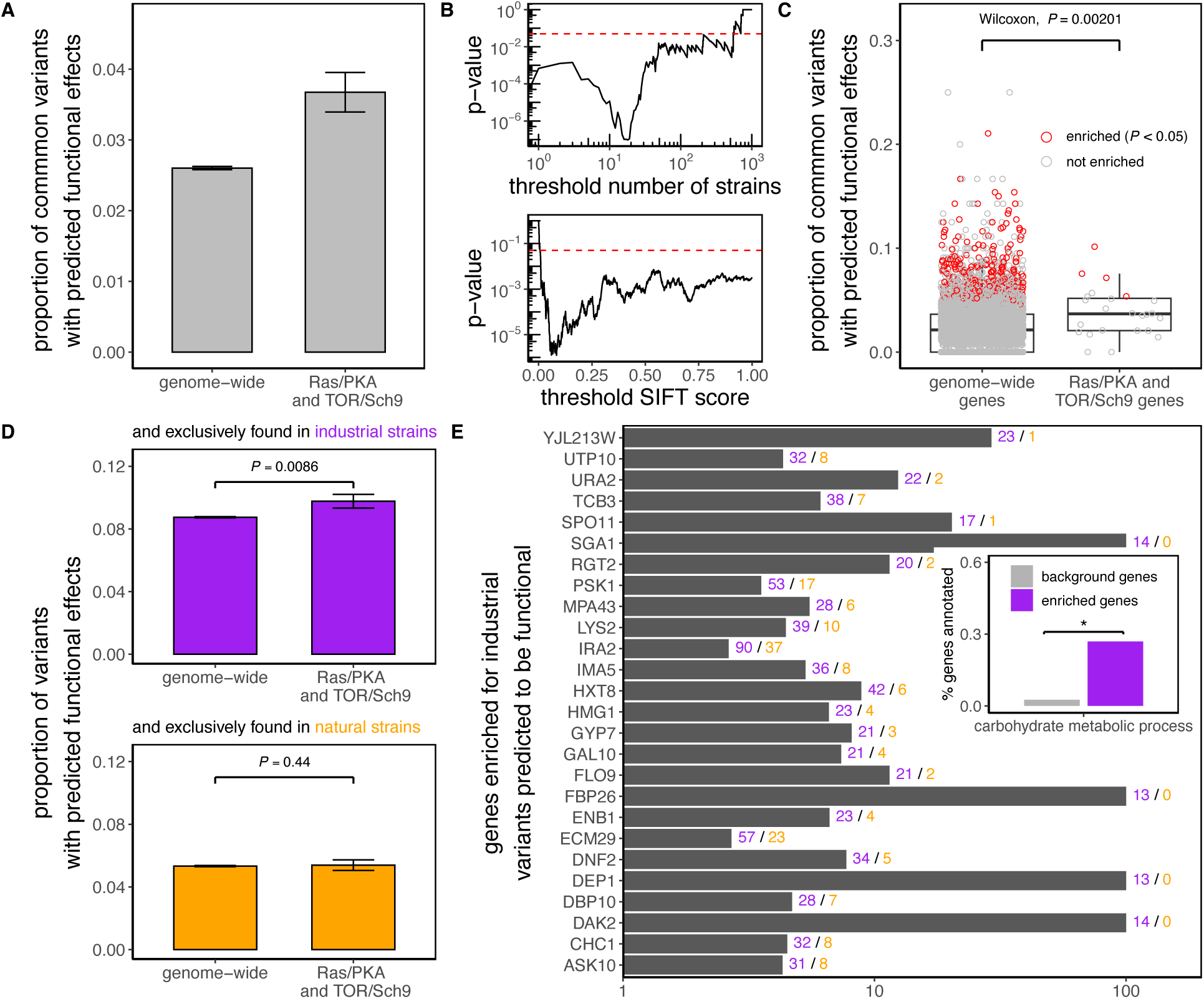
Ras/PKA and TOR/Sch9 genes are genomic hotspots for local adaptation to industrial environments, although other genes likely contribute. (A) Bar plot showing the proportion of non-synonymous variants that are common (>10 strains) and predicted to be functional (SIFT score < 0.05) for RAS/PKA and TOR/Sch9 pathway genes compared to genome-wide genes. Differences were assessed using Fisher’s exact test. (B) The higher proportion of low-SIFT, common non-synonymous variants among RAS/PKA and TOR/Sch9 genes remains robust when varying the minimum number of strains required to define a “common variant” and the maximum SIFT score used to classify a variant as functional. (C) Boxplot showing the proportion of common, functional non-synonymous variants for all 5,359 *S. cerevisiae* genes. RAS/PKA and TOR/Sch9 genes are indicated separately, and generally have higher proportions (Wilcoxon’s test). Individual genes enriched relative to the genomic background are highlighted in red (Fisher’s exact test, *P* < 0.05). (D) Bar plots showing the proportion of non-synonymous variants that are predicted to be functional and exclusive to either industrial (purple) or natural (orange) strains. Variants in RAS/PKA and TOR/Sch9 genes are compared to genome-wide genes. Statistical significance was assessed using Fisher’s exact test (*P* < 0.05). (E) Genes significantly enriched for industrial variants predicted to be functional (FDR-adjusted *P* < 0.05) are shown as a bar plot of odds ratios (industrial vs. natural strains). Counts of variants in industrial and natural strains are indicated to the right of each bar (purple/industrial, orange/natural). Inset shows Gene Ontology (GO) biological process enrichment for the target genes. Only carbohydrate metabolic process is significantly enriched, indicated by an asterisk (*). Bars show the proportion of target genes annotated to this term compared to background genes. In panels A and D bars represent mean ± standard error.

**Supplementary Table S1:** Genes considered in this study.

| Standard name | Systematic name | Pathway |
| --- | --- | --- |
| BCY1 | YIL033C | Ras/PKA |
| CDC25 | YLR310C | Ras/PKA |
| CYR1 | YJL005W | Ras/PKA |
| GPA2 | YER020W | Ras/PKA |
| GPB1 | YOR371C | Ras/PKA |
| GPB2 | YAL056W | Ras/PKA |
| GPR1 | YDL035C | Ras/PKA |
| IRA1 | YBR140C | Ras/PKA |
| IRA2 | YOL081W | Ras/PKA |
| PDE1 | YGL248W | Ras/PKA |
| PDE2 | YOR360C | Ras/PKA |
| RAS1 | YOR101W | Ras/PKA |
| RAS2 | YNL098C | Ras/PKA |
| RGS2 | YOR107W | Ras/PKA |
| TFS1 | YLR178C | Ras/PKA |
| TPK1 | YJL164C | Ras/PKA |
| TPK2 | YPL203W | Ras/PKA |
| TPK3 | YKL166C | Ras/PKA |
| KOG1 | YHR186C | TOR/Sch9 |
| LST8 | YNL006W | TOR/Sch9 |
| SCH9 | YHR205W | TOR/Sch9 |
| TCO89 | YPL180W | TOR/Sch9 |
| TOR1 | YJR066W | TOR/Sch9 |
| TOR2 | YKL203C | TOR/Sch9 |

